# NIPBL-mediated 3D genome folding translates enhancer priming into gene activation and safeguards lineage fidelity during embryonic transitions

**DOI:** 10.64898/2025.12.30.686824

**Authors:** Zhangli Ni, Xiaohui Ma, Stephanie C. Do, Lingling Cheng, Rachel A. Glenn, Thomas Vierbuchen, Alexandros Pertsinidis

## Abstract

Precise gene control by complex regulatory landscapes is fundamental to embryo development, yet the instructive role of 3D genome architecture remains controversial. While acute cohesin depletion completely disrupts genome folding, it yields modest transcriptional impacts, but these findings are often confounded by cohesin’s essential roles in cell division and proliferation. Here, we resolve this discrepancy by decoupling architectural functions from cell-cycle roles using an acute NIPBL degron system. By integrating single-gene imaging with single-cell and bulk multi-omics during mouse pluripotency transitions and germ-layer specification, we show that NIPBL-mediated cohesin function is required for proper *de novo* activation of lineage-specifying genes. Mechanistically, NIPBL translates epigenetic priming into transcriptional outputs by physically bringing distal enhancers and target promoters into proximity. We further uncover a dual regulatory role: an acute requirement for establishing new enhancer–promoter interactions during cell state transitions and a long-term role in safeguarding transcriptional fidelity by preventing ectopic gene de-repression. Our findings demonstrate that NIPBL/cohesin-orchestrated genome folding facilitates the faithful execution of developmental gene expression programs.

**Highlights:**

- Acute NIPBL depletion decouples the architectural functions of cohesin from its essential roles in chromosome segregation and cell cycle progression.
- NIPBL-mediated loop extrusion is required to translate the epigenetic priming of distal enhancers into de novo gene activation during embryonic state transitions.
- NIPBL is a “rate-limiting physical relay” required to bring distal enhancers and target promoters into proximity to initiate transcription.
- 3D genome architecture serves a dual role: enabling acute enhancer–promoter communication and safeguarding long-term lineage fidelity by preventing ectopic gene de-repression.

## Introduction

During early embryonic development, cells must precisely activate and silence specific genes to establish gene expression programs that define lineage identity and guide differentiation^1–3^. Complex regulatory landscapes, comprising multiple *cis*-elements, like enhancers, promoters, silencers, and insulators, over extended genomic distances, are critical for achieving precise gene control in space and time^4–7^, to both maintain transcriptional states and enable rapid reprogramming in response to developmental cues. The 3D genome organization, spanning hierarchical layers from A/B compartments to topologically associated domains (TADs) and chromatin loops, is thought to play important roles, by configuring the physical landscape through which regulatory elements communicate^8–10^.

Cohesin is a key facilitator of genome folding^11,12^, by bringing distal loci into proximity through its loop extrusion activity^13–16^. Barriers to loop extrusion from architectural proteins such as CTCF, help sculpt the main features of 3D genome folding, such as TADs and chromatin loops^11,17,18^. These cohesin-mediated architectures are proposed to contribute to gene regulation by at least two mechanisms: 1) facilitating communication between enhancers and promoters, particularly over large genomic distances^19–25^, by helping bring two loci into proximity; 2) constraining the effects of enhancers into “insulated” regulatory domains and thus selectively activating target genes^12,26–32^. Yet, despite the widespread assumption that 3D architecture underlies transcriptional control, whether cohesin directly contributes to gene regulation remains controversial. Acute cohesin depletion leads to profound structural disruption but surprisingly modest transcriptional effects^11,33^, suggesting that the regulatory roles of cohesin may be highly context-dependent or masked by compensatory mechanisms.

Systematically evaluating cohesin functions in gene regulation, especially over developmental time scales, has been challenging due to the multiple essential roles of cohesin for chromosome segregation and cell division^34,35^. Nipped-B-like protein (NIPBL) is a key cohesin-associated factor that plays an essential role in cohesin loading and loop extrusion^15,16,36^. Haploinsufficiency for NIPBL in humans causes the developmental disorder Cornelia de Lange syndrome (CdLS), and *Nipbl*-heterozygous (*Nipbl*+/-) mice likewise exhibit defects in embryonic development^37–40^. Despite reduced NIPBL dosage, cell division remains largely normal in CdLS patients, *Nipbl*+/− mice, and *Nipbl*-knockdown cell lines^39,41–44^. In addition, mechanistic studies suggest that NIPBL associates with enhancers and modulates promoter activity to influence transcription^45,46^. However, the extent to which NIPBL organizes genome architecture to help establish gene expression programs that maintain and specify cell identities and fates during early embryonic development remains poorly understood.

We develop a degron system targeting NIPBL to disentangle the architectural from the cell-cycle functions of cohesin, and to define its contributions to gene regulation during early mouse embryonic development. The developmental window spanning pre-implantation epiblast to post-implantation gastrulation is fundamental for establishing the mammalian body plan but remains challenging to interrogate *in vivo*^47,48^. Here, we leverage an advanced *in vitro* differentiation systems modeling this developmental progression—from naïve to formative to primed pluripotency, and subsequent germ-layer specification^49,50^—and combine it with single-gene imaging, genomic assays, and single-cell analytics to study whether and how cohesin-mediated genome folding supports establishment and maintenance of new transcriptional programs during cell state transitions.

Our results reveal that NIPBL, and thereby cohesin loop extrusion and cohesin-mediated genome folding, is critical for gene activation during cell state transitions. Loss of cohesin function prevents distal enhancer-promoter interactions, leading to failure to properly induce transcription upregulation, despite correct epigenetic priming and activation of new *cis*-elements. Over extended differentiation time scales, we also observe that cohesin activity is crucial for suppressing ectopic activation of lineage-incompatible genes. Together, these findings suggest two mechanistically and temporally distinct roles for cohesin in gene regulation: an acute requirement for establishing new enhancer–promoter interactions during gene activation, and a longer-term role in safeguarding transcriptional fidelity, likely through domain insulation.

## Results

### Genome conformation is reorganized during the trajectory from naïve embryonic stem cells to definitive endoderm

To dissect the role of the 3D genome topology in cell state maintenance and transitions during early mouse embryonic development, we use an *in vitro* differentiation system that allows for the manipulation of cell fate in a highly controllable manner^49^. In this *in vitro* system, naïve mouse embryonic stem cells (mESCs) can be differentiated into epiblast stem cells (EpiSCs) and further into definitive endoderm (DE) (**Fig. 1A**).

**Figure 1.**
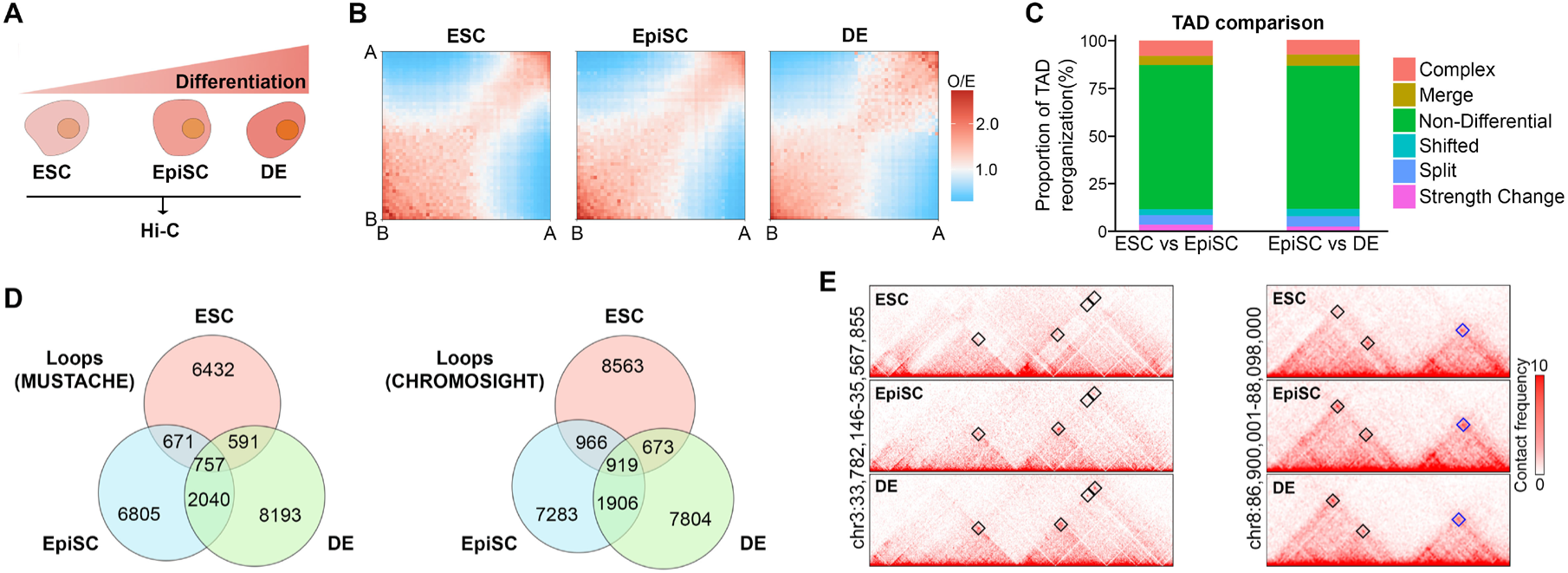
Reorganization of 3D genome architecture during ESC-to–definitive endoderm differentiation. **(A)** Schematic overview of the experimental design. ESC: mouse naïve embryonic stem cell; EpiSC: epiblast stem cell; DE: definitive endoderm. **(B)** Saddle plots showing interactions between A and B compartments in ESCs, EpiSCs, and DE. A and B compartments correspond to transcriptionally active and inactive chromatin domains, respectively. **(C)** TAD boundary changes between related cell types, comparing ESCs with EpiSCs and EpiSCs with DE. Colors indicate different categories of TAD boundary changes, defined based on the differential boundary score and spatial relationships between boundaries: non-differential (unchanged), shifted (boundary position shift within a local window), merged (loss of an intervening boundary), split (emergence of a new boundary), strength change (altered boundary insulation strength), and complex (combinations of multiple boundary changes). **(D)** Venn diagram illustrating the overlap of identified chromatin loops by Mustache and Chromosight loop callers among ESCs, EpiSCs, and DE. Numbers indicate the number of loops in each group. **(E)** Representative Hi-C contact maps in ESCs, EpiSCs, and DE. Black boxes indicate differential loops, and blue boxes mark common loops. Hi-C libraries were generated in parallel with dTAG-treated samples, but only the DMSO control datasets are shown in this figure.

To understand 3D genome organization and its dynamic changes among the three cell types, we perform high-resolution Hi-C using a restriction enzyme cocktail with high cutting frequency. Hi-C data shows high quality, with >89.9% mapping rate, >80% valid interaction pairs, and >58% long-range (>20 kb) intra-chromosomal contacts (**Supp Table 1**). Based on these Hi-C data, we analyze 3D chromatin organization at multiple scales, including A/B compartments, TADs, and chromatin loops. The strength of A/B compartmentalization first decreases during the transition from the naïve pluripotent state (ESCs) to the primed pluripotent state (EpiSCs) and then increases again during the transition to definitive endoderm (**Fig. 1B**). Analysis of TAD boundary changes between related cell types shows that most boundaries are conserved, with 76% remaining unchanged between ESCs and EpiSCs, and 74% remaining unchanged between EpiSCs and DE (**Fig. 1C**). However, at the chromatin loop level, analysis with two loop calling algorithms reveals that 65%-77% of loops were unique to each cell type (**Fig. 1D-E**). Taken together, these results indicate that the 3D genome conformation exhibits cell type-specific organization and undergoes reorganization as naïve embryonic stem cells differentiate into pluripotent epiblast cells and then into definitive endoderm.

### Depletion of NIPBL disrupts 3D genome architecture in naïve embryonic stem cells, epiblast stem cells, and definitive endoderm

To establish experimental systems in which 3D genome organization can be modulated, we focus on cohesin, a major genome architectural factor. We generate knock-in mouse ESCs where the endogenous cohesin subunit RAD21 can be acutely depleted using the dTAG-inducible degron system (**Supp Fig. 1A-B**)^51^. However, as reported previously^44^, we observe that RAD21 depletion results in abnormal chromosome segregation during cell division, characterized by lagging chromosomes and delayed sister chromatid separation (**Supp Fig. 1C**). Studying roles of 3D genome topology in cell state maintenance and transitions during developmental time scales requires perturbing 3D genome organization without affecting cell division. As an alternative to perturbing cohesin complex activity, we focus on NIPBL, a cohesin-associated factor that is required for cohesin loading and processive loop extrusion. We homozygously tag the endogenous NIPBL with dTAG-inducible degron in mESCs (**Fig. 2A**). Time-course analysis shows that NIPBL is acutely degraded after dTAG treatment, reaching maximum depletion at 4 h (**Fig. 2B-C** and **Supp Fig. 1D**). Contrary to the RAD21 degron, after NIPBL depletion, most cells (96%) still exhibit sharp and rapid chromosome segregation comparable to that of control cells (**Supp Fig. 1E**).

**Figure 2.**
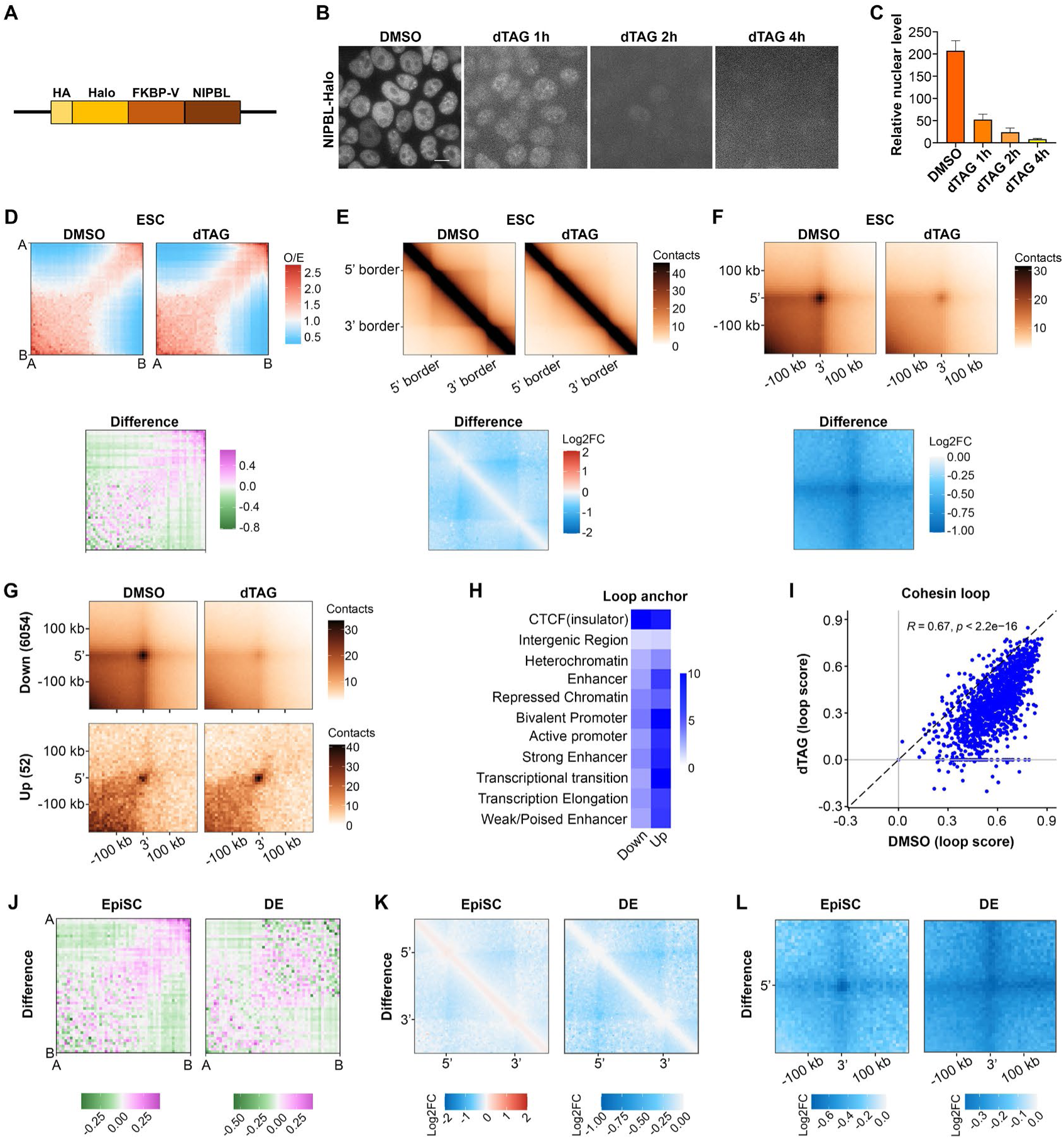
NIPBL depletion alters 3D genome architecture in mESCs, EpiSCs, and definitive endoderm. **(A)** Schematic overview of endogenous editing NIPBL for degradation using the dTAG degron system. **(B)** Representative fluorescence images showing time-dependent degradation of NIPBL-Halo in mESCs. NIPBL–Halo was labeled with HaloTag ligand prior to imaging. Scale bar = 10 µm. **(C)** Bar plot showing relative nuclear levels of NIPBL–Halo quantified from fluorescence images. **(D)** Upper saddle plots showing interactions between A and B compartments in DMSO-treated and dTAG-treated mESCs. Lower differential saddle plot illustrating changes in compartment strength between the two conditions. Positive values indicate stronger compartment interactions in dTAG-treated cells relative to DMSO controls. **(E)** Upper panels show aggregate TAD analysis (ATA) displaying all TAD contacts in DMSO-treated and dTAG-treated mESCs. Lower panel shows differential ATAs illustrating changes in TAD contact strength between the two conditions. Negative values indicate weaker TAD contact strength in dTAG-treated cells relative to DMSO controls. **(F)** Upper panels show aggregate loop analysis (APA) displaying all loop contacts in DMSO-treated and dTAG-treated mESCs. Lower panel shows differential APAs illustrating changes in loop contact strength between the two conditions. Negative values indicate weaker loop contact strength in dTAG-treated cells relative to DMSO controls. **(G)** APAs showing loops classified as downregulated (Down) or upregulated (Up) by Mustache following dTAG treatment. **(H)** Enrichment analysis of ChromHMM chromatin-state at loop anchors associated with upregulated or downregulated loops. **(I)** Scatter plot of loop scores detected by Chromosight for cohesin loops in DMSO-treated and dTAG-treated mESCs. **(J)** Differential saddle plots illustrating changes in compartment strength between the two conditions (DMSO and dTAG) in EpiSC and DE. Positive values indicate stronger compartment interactions in dTAG-treated cells relative to DMSO controls. **(K)** Differential ATAs illustrating changes in TAD contact strength between the two conditions (DMSO and dTAG) in EpiSC and DE. Negative values indicate weaker TAD contact strength in dTAG-treated cells relative to DMSO controls. **(L)** Differential ATAs illustrating changes in TAD contact strength between the two conditions (DMSO and dTAG) in EpiSC and DE. Negative values indicate weaker loop contact strength in dTAG-treated cells relative to DMSO controls. In Hi-C experiments, acute NIPBL depletion is performed for 4 h in mESCs and EpiSCs, while the NIPBL degradation strategy used in DE is shown in Fig. 5I.

Subsequently, we investigate whether NIPBL is involved in the modulation of genome conformation in the three cell types using Hi-C analysis. At scales >2 Mb, genome-wide aggregate analysis reveals increased A-A and B-B compartment interactions and decreased A-B interactions, indicating that genome compartmentalization is strengthened upon acute NIPBL depletion in mESCs (**Fig. 2D** and **Supp Fig. 2A**), consistent with effects of cohesin depletion on compartment interactions^11,17,33^. In contrast, at smaller scales (<2 Mb), NIPBL depletion weakens contacts within TADs and chromatin loops in mESCs (**Fig. 2E-F**). Chromatin loops can be broadly categorized into structural loops, formed by cohesin halted at convergent CTCF sites, and cis-regulatory element (CRE) loops, which are generally more dynamic and often associated with enhancer-promoter communication^11,52–57^. To further characterize the effect of NIPBL depletion on the different loop types, we analyze changes in loop strength. 6054 out of 8451 (72% of all loops) are significantly down-regulated and 52 (0.6%) are up-regulated (**Fig. 2G**). Both up- and down-regulated loop anchors are significantly enriched for CTCF sites (**Fig. 2H**), consistent with CTCF’s role in organizing chromatin loops along with cohesin^17^. Furthermore, the enrichment of ChromHMM chromatin-state at loop anchors indicates that two groups of loop anchors are enriched at promoters and enhancers, with stronger enrichment at upregulated loops. Correlation analysis of cohesin loop strength between DMSO and dTAG conditions reveals that most (>90%) cohesin loops are weakened upon NIPBL depletion in mESCs (**Fig. 2I**). These results demonstrate that NIPBL modulates 3D genome organization by mediating cohesin loop extrusion in mESCs and may directly contribute to the formation of CRE-associated loops.

Consistent with the results in ESCs, Hi-C experiments after NIPBL depletion in EpiSCs and DE reveal comparable changes of genome conformation, including strengthened compartmentalization and weakened intra-TADs and chromatin loop contacts (**Fig. 2J-L** and **Supp Fig. 2B-C**). Overall, these findings highlight the conserved role of NIPBL in modulating 3D genome architecture across the three cell types.

### Naïve embryonic stem cell state shows robustness to NIPBL depletion, with only a modest differentiation phenotype that is reversible upon NIPBL recovery

Since NIPBL depletion has minimal effects on cell division but strongly alters 3D genome architecture, we wonder whether long-term NIPBL depletion affects the maintenance of naïve embryonic stem cell state. By examining the morphology of stem cell colonies, we find that most cells still grow in colonies after 6 days of NIPBL depletion, although slightly more cells are observed outside the colonies compared with the control group (**Fig. 3A** and **Supp Fig. 3**). By qPCR analysis to detect the expression of pluripotency genes, we also observe that the expression of *Klf4* and *Tbx3* was significantly reduced after NIPBL depletion, whereas *Sox2* expression remained unchanged (**Fig. 3B**).

**Figure 3.**
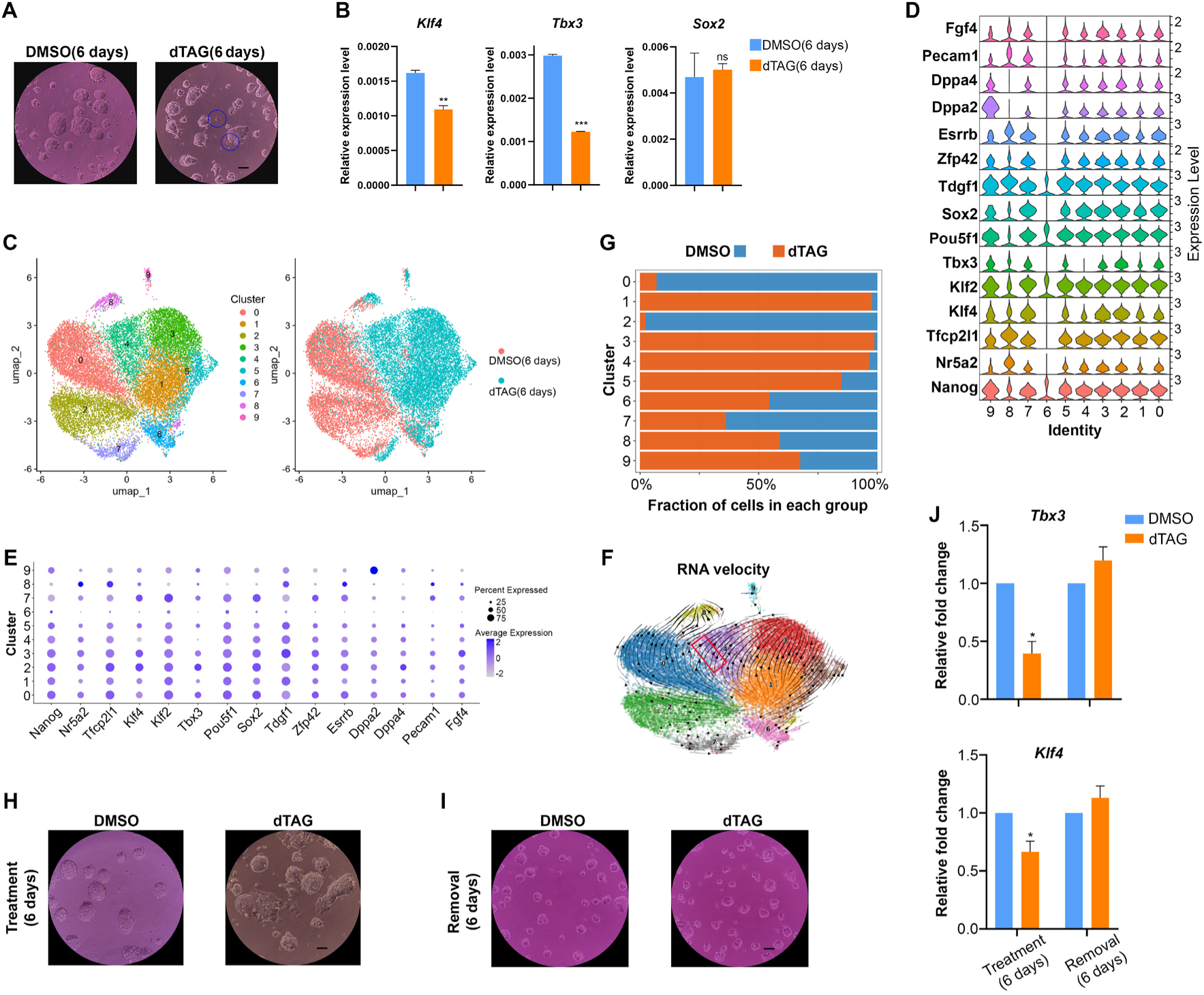
ESC fate maintenance is robust to NIPBL depletion, and NIPBL recovery rescues the modest differentiation phenotype. **(A)** Brightfield images showing mESC colonies after treatment with DMSO or dTAG for 6 days. Blue circles highlight cells that grow out of colonies in the dTAG-treated sample. Scale bar = 100 µm. **(B)** qPCR analysis of *Klf4*, *Tbx3*, and *Sox2* expression in mESCs after treatment with DMSO or dTAG for 6 days. Values were normalized to *Gapdh* expression. Data are presented as mean ± SEM. Two-tailed unpaired Student’s t-test; **P < 0.01, ***P < 0.001, ns, not significant. **(C)** Left: UMAP plot showing cell clusters from the integrated datasets of mESCs treated with DMSO or dTAG for 6 days. Right: UMAP plot showing DMSO-treated mESCs in red and dTAG-treated mESCs in blue. **(D)** Violin plots illustrating the expression levels of pluripotency genes in each cell cluster. **(E)** Dot plots showing the average expression levels (color) and the percentage of expressing cells (dot size) for pluripotency genes across cell clusters. **(F)** Fraction of DMSO- and dTAG-treated mESCs in each cluster. **(G)** RNA velocity analysis. Arrows indicate the direction of differentiation. **(H)** Brightfield images showing mESC colonies after treatment with DMSO or dTAG for 6 days. Scale bar = 100 µm. **(I)** Brightfield images showing mESC colonies 6 days after DMSO or dTAG removal. Scale bar = 100 µm. **(J)** qPCR analysis of Klf4 and Tbx3 expression in mESCs treated as in panels (**H**) and (**I**). Values were normalized to *Gapdh* expression, and expression levels in the dTAG groups were further normalized to the DMSO control. *P < 0.05.

To better understand cell heterogeneity after NIPBL depletion, we analyze DMSO- and dTAG-treated mESCs by single-cell RNA sequencing (scRNA-seq). After quality control and integration of the two datasets (DMSO and dTAG), we identify 10 distinct cell clusters, visualized by UMAP analysis in **Fig. 3C**. Cells in different cell cycle phases (G1, S, and G2/M) are evenly distributed across clusters (**Supp Fig. 4**), indicating that the clustering is independent of cell cycle effects.

**Figure 4.**
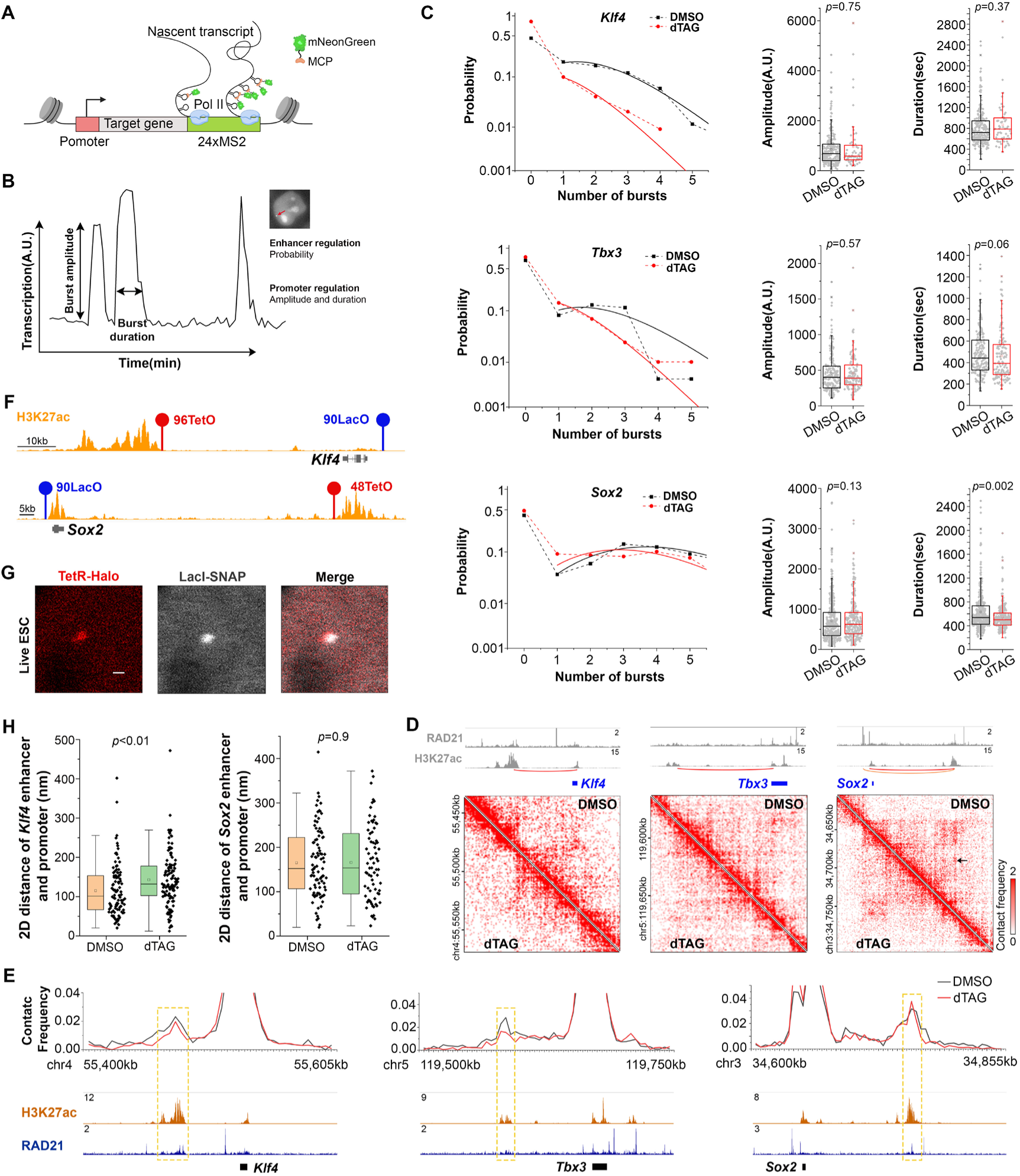
NIPBL regulates transcriptional bursting of *Klf4* and *Tbx3*, but not *Sox2*, by modulating enhancer–promoter spatial proximity in ESCs. **(A)** Schematic of live-cell real-time imaging of nascent transcription using the MS2/MCP system. Created with BioRender.com. **(B)** Schematic illustrating transcriptional bursting. **(C)** Burst probability, amplitude, and duration of *Klf4*, *Tbx3*, and *Sox2* in DMSO- and dTAG-treated cells. Each gray dot in the box plots of burst amplitude and duration represents one burst. Data are from two independent experiments. **(D)** Hi-C heatmaps showing chromatin contacts in DMSO-treated (top right) and dTAG-treated (bottom left) mESCs around the *Klf4*, *Tbx3*, and *Sox2* loci. The top panels show ChIP-seq signal tracks for RAD21 and H3K27ac in the same regions, using published datasets. Arrow indicates a stripe structure between Sox2 and the SCR. **(E)** Virtual 4C profiles quantifying contact frequencies in DMSO-treated and dTAG-treated mESCs around the *Klf4*, *Tbx3*, and *Sox2* loci. Distal enhancers are highlighted by yellow dashed boxes. The bottom panels show ChIP-seq signal tracks for H3K27ac and RAD21 in the same regions, using published datasets. **(F)** Positions of LacO and TetO array insertions are indicated on H3K27ac ChIP-seq signal tracks at the *Klf4* and *Sox2* loci. LacO arrays are marked in blue, and TetO arrays are marked in red. **(G)** Representative confocal images showing TetR-Halo and LacI-SNAP foci in live mESCs. TetR-Halo and LacI-SNAP were labeled with Halo-tag and SNAP-tag ligands and are shown in red and grey, respectively. Scale bar = 400 nm. **(H)** 2D distance measurements between the enhancers and promoters of *Klf4* and *Sox2*, labeled with TetO/TetR-Halo and LacO/LacI-SNAP, respectively. Each black dot to the right of the box plots represents one data point. Data are from two independent experiments.

We next characterize these cell clusters according to the expression of pluripotency genes and find that 8 major clusters highly express key pluripotency factors, including *Sox2*, *Pou5f1*, and *Nanog*, except for clusters 6 and 8 (**Fig. 3D-E**). None of the top 10 differentially expressed genes identified by scRNA-seq are genes that are important for pluripotency (**Supp Fig. 5**). Nonetheless, clusters 1, 4, and 5, which mainly consist of dTAG-treated cells, show lower expression of naïve pluripotency genes, such as *Klf4* and *Tbx3*, compared with clusters 0 and 2, which are primarily composed of DMSO-treated cells (**Fig. 3D-F**). RNA velocity analysis further reveals a potential differentiation trajectory from cluster 0 to cluster 4 (**Fig. 3F**). In addition, the fraction of cells in the differentiated clusters 6 and 8 are slightly higher in the dTAG- than in the DMSO-treated sample (**Fig. 3G**). These results indicate that although the naïve embryonic stem cell fate remains largely robust after NIPBL depletion, the expression of naïve pluripotency genes is reduced.

**Figure 5.**
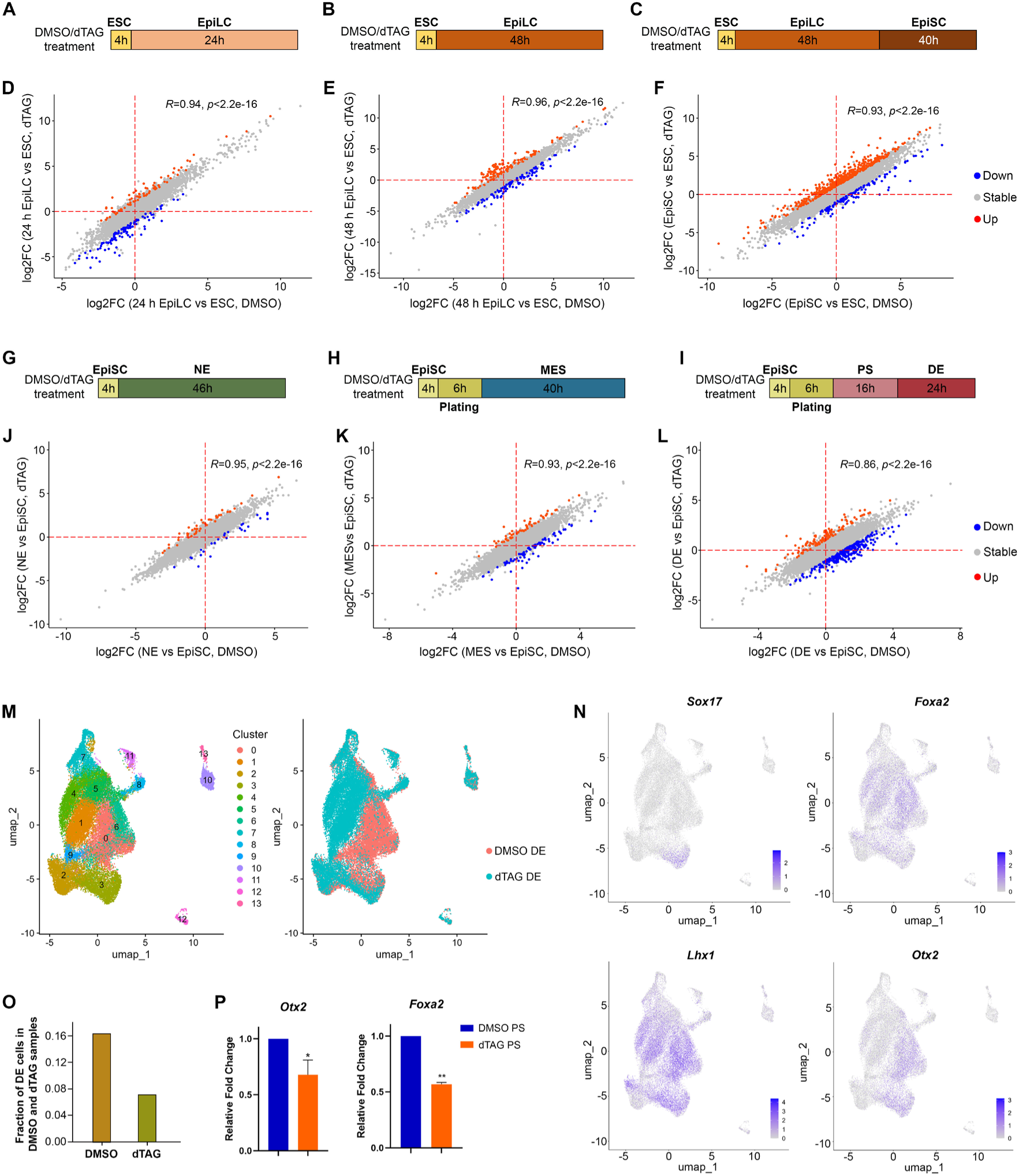
NIPBL is required for gene activation during pluripotent cell fate transitions and definitive endoderm lineage specification. **(A)(B)(C)** Schematic overview of EpiLC and EpiSC differentiation and the timing of DMSO or dTAG treatment during differentiation. **(D)(E)(F)** Correlation analysis of activated and deactivated genes during the differentiation of mESCs into EpiLCs and EpiSCs between DMSO- and dTAG-treated samples. Downregulated genes after dTAG treatment are marked in blue, and upregulated genes are marked in red. P < 0.01, |fold change| ≥ 2. **(G)(H)(I)** Schematic overview of three germ layer differentiation and the timing of DMSO or dTAG treatment during differentiation. **(J)(K)(L)** Correlation analysis of activated and deactivated genes during the differentiation of EpiSCs into neuroectoderm (NE), mesoderm (MES), and definitive endoderm (DE) between DMSO-and dTAG-treated samples. Downregulated genes after dTAG treatment are marked in blue, and upregulated genes are marked in red. P < 0.01, |fold change| ≥ 2. PS: primitive streak. **(M)** Left: UMAP plot showing cell clusters from the integrated datasets of DE cells treated with DMSO or dTAG, following the treatment scheme shown in (**I**). Right: UMAP plot showing DMSO-treated DE cells in red and dTAG-treated DE cells in blue. **(N)** Feature plots showing the expression of *Sox17*, *Foxa2*, *Lhx1*, and *Otx2* across different clusters. **(O)** Fraction of DE cells in DMSO- and dTAG-treated samples. **(P)** qPCR analysis of *Otx2* and *Foxa2* expression in PS treated as shown in (**I**). Values were normalized to *Gapdh* expression, and expression levels in the dTAG groups were further normalized to the DMSO control. *P < 0.05. **P < 0.01.

Interestingly, cluster 9 is clearly separated from the other clusters on the UMAP plot, despite being in a pluripotent state. Gene expression analysis reveals that this cluster expresses makers of 2-cell embryo (**Supp Fig. 6**). This observation is consistent with previous reports showing the presence of a small population of 2-cell-like cells within naïve embryonic stem cell cultures^58^, suggesting that cluster 9 represents a 2-cell-like subpopulation in the pluripotent state.

**Figure 6.**
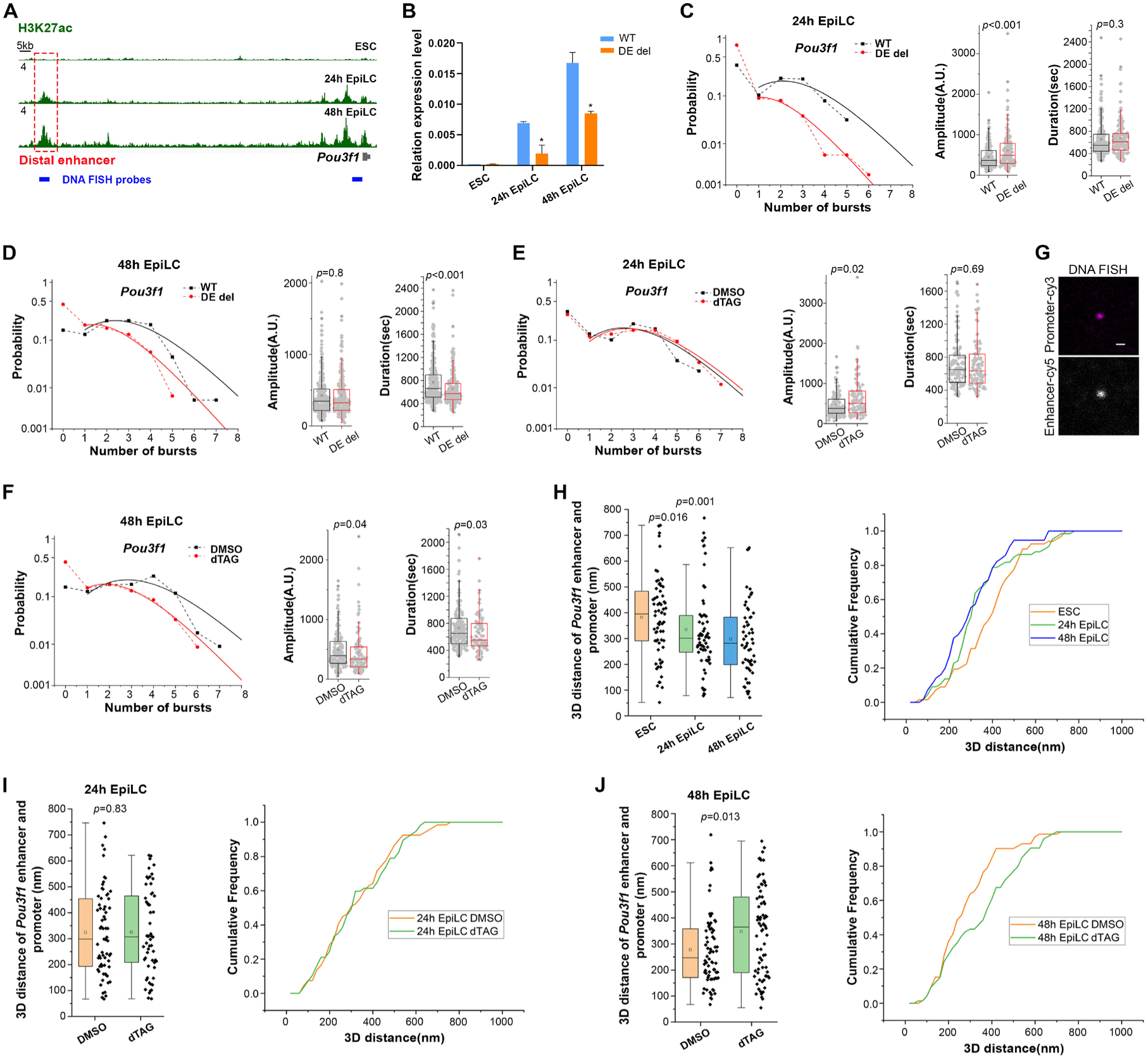
NIPBL mediates spatial interactions between distal enhancer and the *Pou3f1* promoter to enable full activation in EpiLCs. **(A)** ChIP-seq signal tracks of H3K27ac around the *Pou3f1* locus during the differentiation of mESCs into EpiLCs, using published datasets. Distal enhancer is highlighted by red dashed box. **(B)** qPCR analysis of *Pou3f1* expression during the differentiation of mESCs into EpiLCs before and after distal enhancer deletion. Values were normalized to *Gapdh* expression. Data are presented as mean ± SEM. Two-tailed unpaired Student’s t-test; *P < 0.05. **(C)(D)** Burst probability, amplitude, and duration of *Pou3f1* in 24 h and 48 h EpiLCs before and after distal enhancer deletion. Each gray dot in the box plots of burst amplitude and duration represents one burst. Data are from three independent experiments. **(E)(F)** Burst probability, amplitude, and duration of *Pou3f1* in 24 h and 48 h EpiLCs treated with DMSO and dTAG, following the treatment scheme shown in Fig. 5A and B. Each gray dot in the box plots of burst amplitude and duration represents one burst. Data are from two independent experiments. **(G)** Representative confocal images showing *Pou3f1* enhancer–Cy5 and promoter–Cy3 foci in OligoPAINT DNA FISH samples. Scale bar = 400 nm. **(H)** 3D distance measurements between the enhancers and promoters of *Pou3f1* during the differentiation of mESCs into EpiLCs, labeled with Cy5 and Cy3, respectively. Each black dot to the right of the box plots represents one data point. **(I)(J)** 3D distance measurements between the enhancers and promoters of *Pou3f1* in 24 h and 48 h EpiLCs treated with DMSO and dTAG, following the treatment scheme shown in Fig. 5A and B. Each black dot to the right of the box plots represents one data point.

Based on the observation that NIPBL depletion results in slightly more cells growing out of ESC colonies and reduced expression levels of *Klf4* and *Tbx3*, we ask whether removal of dTAG could reverse this phenotype. Removal of dTAG for 6 days, which restores NIPBL expression, rescues the differentiation-like phenotype and restores the expression levels of Klf4 and Tbx3 to levels comparable to those in DMSO-treated samples (**Fig. 3H-J** and **Supp Fig. 7**).

**Figure 7.**
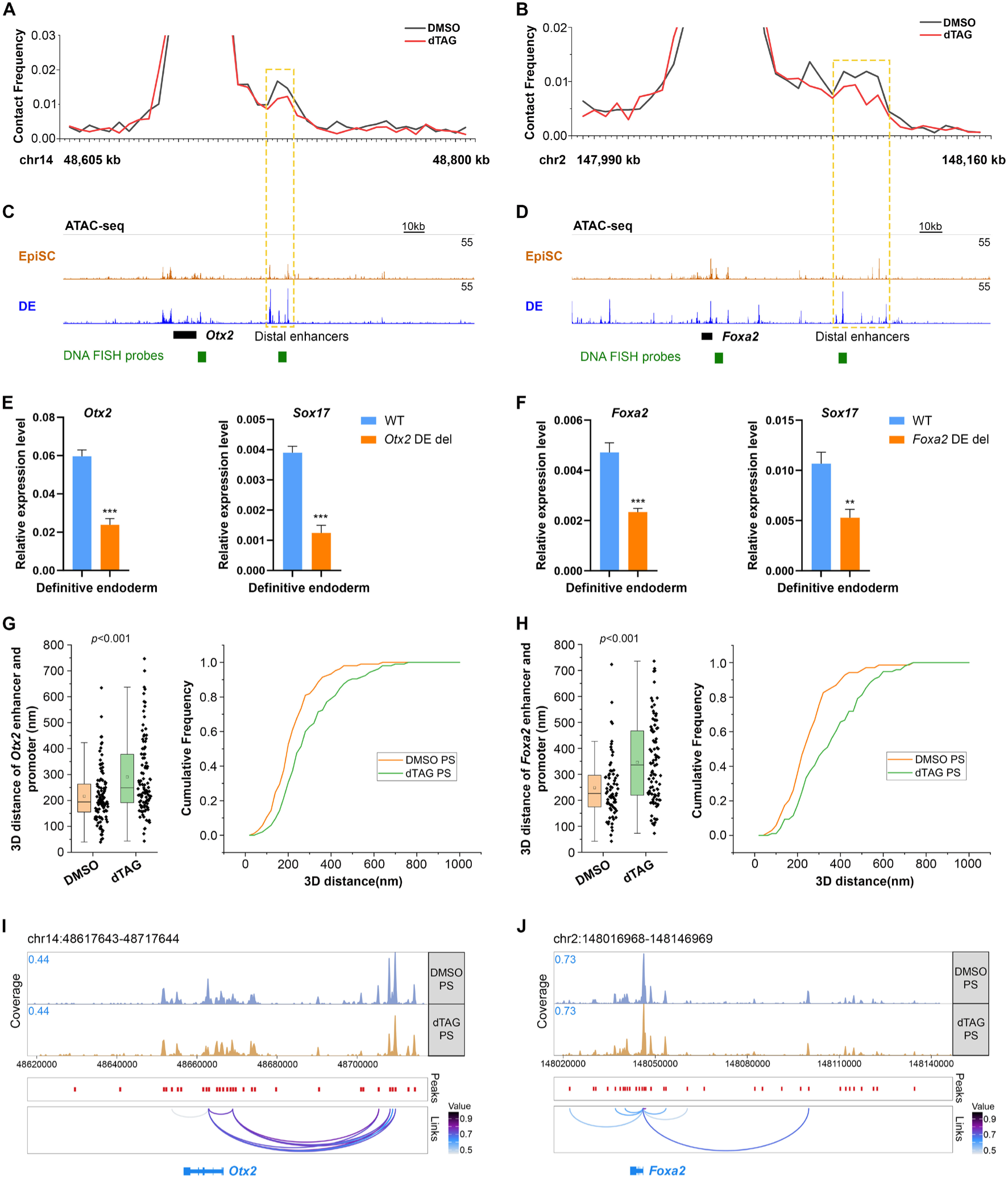
NIPBL depletion disrupts enhancer–promoter interactions and proximity at the *Otx2* and *Foxa2* loci during definitive endoderm differentiation without altering chromatin accessibility. **(A)(B)** Virtual 4C profiles quantifying contact frequencies in DMSO-treated and dTAG-treated definitive endoderm around the *Otx2* and *Foxa2* loci. **(C)(D)** ATAC-seq signal tracks in the same regions as shown in (A) and (B), using published datasets. Distal enhancers are highlighted by yellow dashed boxes. **(E)** qPCR analysis of *Otx2* and *Sox17* expression in definitive endoderm before and after deletion of the *Otx2* distal enhancer. Values were normalized to *Gapdh* expression. Data are presented as mean ± SEM. Two-tailed unpaired Student’s t-test; ***P < 0.001. **(F)(G)** 3D distance measurements between the enhancers and promoters of *Otx2* and *Foxa2* in primitive streak treated with DMSO and dTAG, following the treatment scheme shown in Fig. 5I. Each black dot to the right of the box plots represents one data point. **(H)(I)** Browser tracks of peak-to-gene links derived from scMultiomics data show comparable chromatin accessibility between DMSO- and dTAG-treated PS at the *Otx2* and *Foxa2* loci. PS: primitive streak.

Collectively, these findings demonstrate that the naïve embryonic stem cell fate exhibits robustness to NIPBL depletion, and that the modest differentiation phenotype is reversible upon NIPBL recovery.

### NIPBL controls transcriptional bursting kinetics of affected pluripotency genes by modulating enhancer-promoter spatial encounters in ESCs

To further investigate the mechanism by which NIPBL regulates naïve pluripotency genes in mESCs, we employ live-cell real-time imaging of nascent transcription using the MS2/MS2 coat protein (MCP) system^59^ to evaluate the effects of acute perturbations on nascent transcription kinetics (**Fig. 4A**). The temporal dynamics of these genes show On-Off bursting behaviors^60,61^. We thus analyze burst probability, burst amplitude, and burst duration (**Fig. 4B**) to define transcriptional activity in live cells. Changes in burst amplitude and duration are proposed to be associated with promoter regulation, whereas burst probability—reflecting the fraction of promoters in a state permissive to bursting as well as the frequency of bursts—is associated with enhancer regulation^62–64^. Acute NIPBL depletion (4 h) results in more than 4-fold and more than 2-fold decrease in burst probability for *Klf4* and *Tbx3*, respectively, while *Sox2* exhibits a milder change (**Fig. 4C**, left). Meanwhile, burst amplitude and burst duration are only modestly affected, with a ∼10–15% median shift, or remain unaffected (**Fig. 4C**, right two columns). These results indicate that NIBPL predominantly modulates enhancer-controlled transcriptional burst kinetics, and that NIPBL depletion has a stronger effect on burst probability of *Klf4* and *Tbx3* than *Sox2*.

The above results describe NIPBL’s functions in asynchronous interphase cells. However, cohesin and CTCF appear to play a more pivotal role in driving post-mitotic chromatin reconfiguration and enabling gene reactivation during mitotic exit^65,66^. To further examine the impact of NIPBL depletion on gene reactivation, we synchronize mESCs in mitosis and track transcriptional bursting after release (**Supp Fig. 8A**). Consistently, NIPBL depletion causes a more severe defect in burst probability (∼3-fold decrease) during *Sox2* reactivation, whereas *Klf4* shows a response comparable to its interphase nascent transcription (**Supp Fig. 8B-C**). Notably, once *Sox2* becomes fully reactivated, the effect of NIPBL depletion diminishes (**Supp Fig. 8D-E**). These results suggest that *Sox2* reactivation during cell division is more dependent on NIPBL than the maintenance of its expression afterwards.

**Figure 8.**
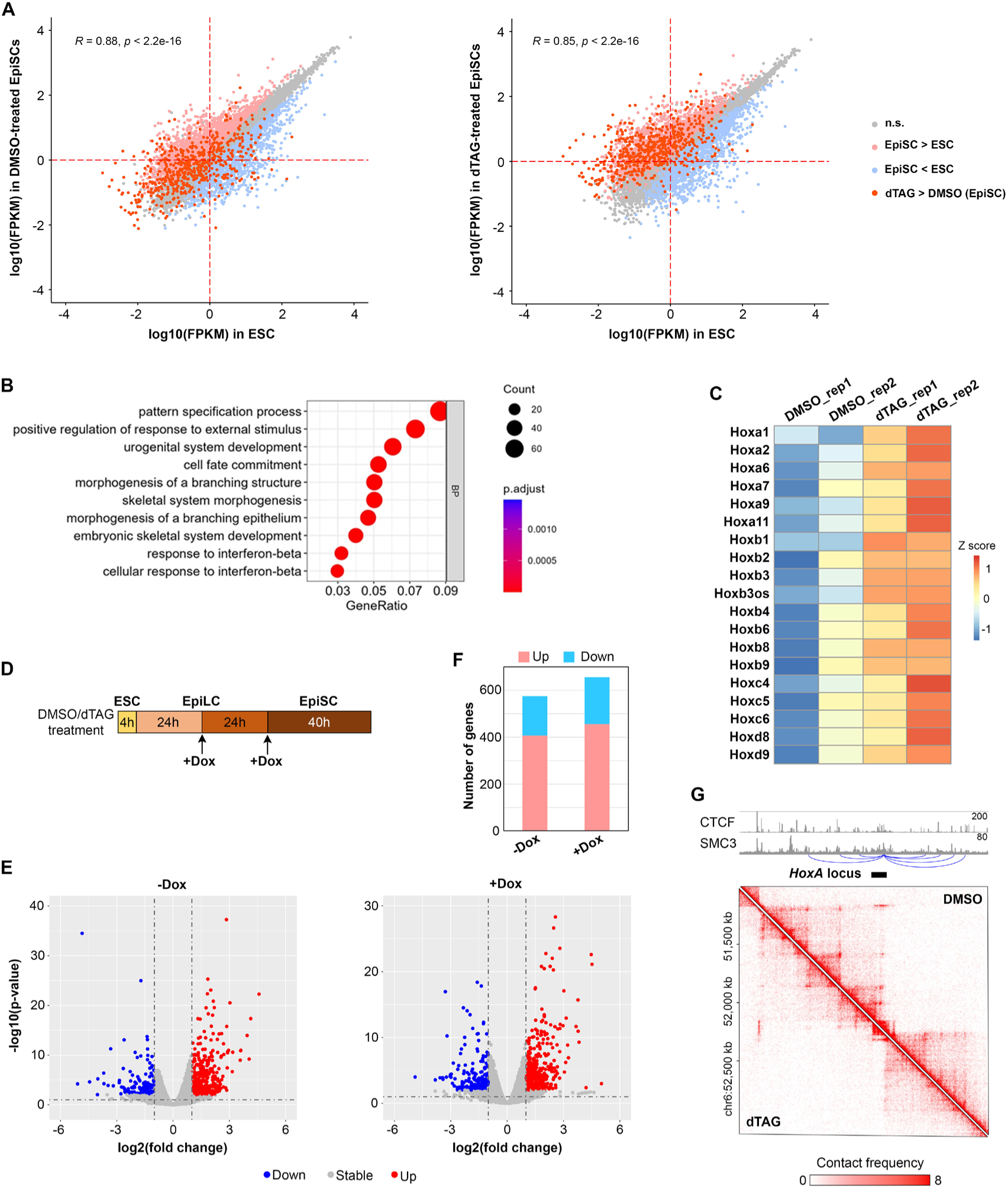
Long-term NIPBL depletion leads to derepression of lineage-inappropriate genes during the ESC-to-EpiSC transition. **(A)** Correlation plot of gene expression levels (log₁₀ FPKM) between ESCs and DMSO- or dTAG-treated EpiSCs from an independent biological replicate. Upregulated and downregulated genes in EpiSCs are shown in pink and light blue, respectively, while genes without significant changes are shown in gray. Genes upregulated upon NIPBL depletion in EpiSCs are shown in red. P < 0.01; |fold change| ≥ 2. **(B)** Top GO terms enriched in genes upregulated upon NIPBL depletion in EpiSCs. **(C)** Heatmap showing expression of *Hox* cluster genes in DMSO- or dTAG-treated EpiSCs from two independent biological replicates. **(D)** Schematic showing the timing of DMSO or dTAG treatment during EpiSC differentiation. Doxycycline is added to overexpress DNMT3B in 24 h and 48 h EpiLCs. **(E)** Volcano plots showing differentially expressed genes between DMSO- and dTAG-treated EpiSCs without (-Dox) or with (+Dox) DNMT3B overexpression. Upregulated and downregulated genes are indicated in red and blue, respectively. P < 0.01, |fold change| >= 2. **(F)** Bar graph showing the number of upregulated and downregulated genes without (−Dox) or with (+Dox) DNMT3B overexpression. **(G)** Snapshots of Hi-C maps showing chromatin contacts in DMSO-treated (top right) and dTAG-treated (bottom left) EpiSC around the *HoxA* cluster. Top panels show ChIP-seq signal tracks for CTCF and SMC3 at the same genomic regions, using published datasets from E6.5 mouse embryos.

The live-cell transcription imaging results suggest that NIPBL may control bursting by directly modulating promoter-enhancer interactions. To test this hypothesis, we analyze our Hi-C data in mESCs and quantify promoter-enhancer contact frequencies using virtual 4C analysis. As shown in **Fig. 4D-E**, acute NIPBL depletion in mESCs weakens interactions between the promoters and previously characterized distal enhancers of *Klf4* and *Tbx3*^67,68^. In contrast, the interactions between the *Sox2* promoter and the distal *Sox2* control region (SCR)^69^, are not significantly affected. However, some structural changes at the *Sox2* locus are evident, such as disruption of cohesin loops marked by RAD21 signal, between ∼10 kb upstream of *Sox2* and the SCR, as well as a stripe structure between *Sox2* and SCR. These results indicate that direct enhancer-promoter contacts mediated by NIPBL (rather than structural loops), are critical for gene expression.

To further confirm the effect of acute NIPBL depletion on promoter-enhancer interactions, we generate knock-in cell lines with DNA tags to investigate changes in spatial promoter-enhancer proximity in live cells. We integrate LacO and TetO arrays at the promoters and enhancers, respectively, of *Klf4* and *Sox2* (**Fig. 4F**), and visualize the two loci using LacI-SNAP and TetR-Halo fusion proteins and fluorescent Halo and SNAP ligands (**Fig. 4G**). Consistent with the Hi-C data, 2D distance measurements reveal that acute NIPBL depletion significantly increases the spatial distance between the promoter and enhancer of *Klf4*, but not *Sox2* (**Fig. 4H**).

Taken together, these findings demonstrate that NIPBL controls the transcriptional bursting kinetics of *Klf4* and *Tbx3*, but not *Sox2*, in mESCs, by increasing enhancer-promoter spatial proximity.

### NIPBL function ensures activation of genes important for lineage specification and cell differentiation

Considering that 3D genome architecture undergoes dynamic reorganization during early embryonic development, we wonder whether and how NIPBL and cohesin functions contribute to cell state transitions and transcriptional reprogramming. First, we focus on the role of NIPBL in the differentiation of naïve ESCs into formative (EpiLCs) and subsequently into primed pluripotent cells (EpiSCs) (**Fig. 5A-C**). We perform bulk RNA-seq to characterize gene activation and deactivation during differentiation and to examine transcriptional changes upon NIPBL depletion. Despite the high correlation (R > 0.9) between the gene activation and inactivation profiles of dTAG-treated and DMSO-treated cells, NIPBL depletion leads to differential expression (|fold change| ≥ 2, p-value < 0.01) of hundreds of genes in EpiLCs and about a thousand in EpiSCs (**Fig. 5D-F** and **Supp Fig. 9**). We observe temporally distinct effects of NIPBL depletion. Early in the EpiLC transition (24 h), NIPBL depletion accelerates the downregulation of (naïve) pluripotency genes (such as *Nr5a2*, *Nanog*, *Esrrb*), consistent with the acute effects of NIPBL depletion on these genes in the ESC state. At later stages (48 h) of the EpiLC transition, as well as in EpiSCs, many activated genes, including *Pou3f1* and *Dnmt3b*, fail to reach appropriate expression levels upon NIPBL depletion. These results demonstrate that, although NIPBL depletion does not strictly prevent these cell state transitions, it results in incomplete gene activation.

**Figure 9.**
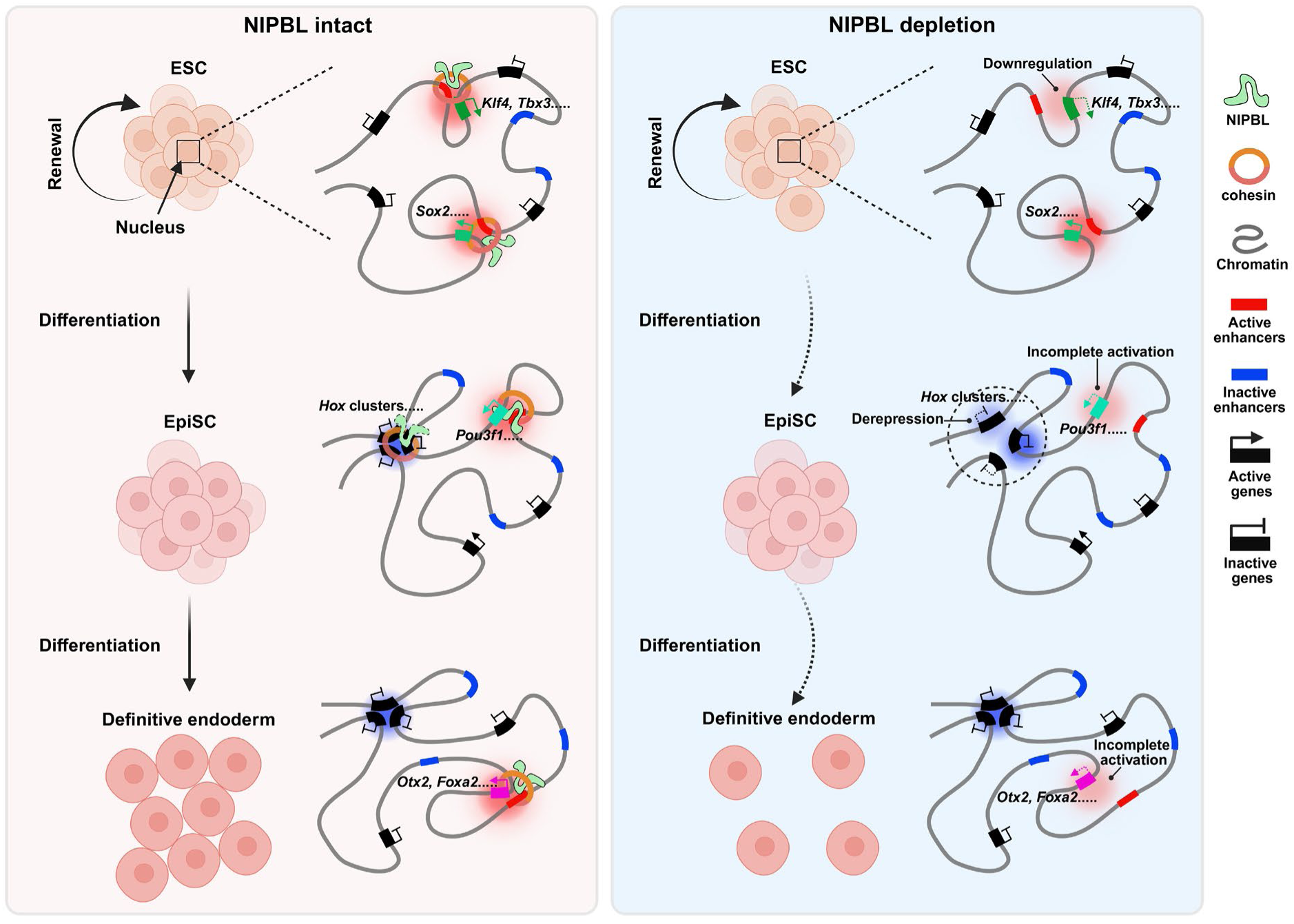
Schematic models illustrating the role of NIPBL, and thereby cohesin-mediated genome folding, in enhancer–promoter interactions during cell fate transitions from naïve pluripotency to definitive endoderm. NIPBL facilitates gene activation by promoting enhancer-promoter proximity during cell fate transitions, whereas long-term NIPBL depletion leads to derepression of lineage-inappropriate genes, likely through altered genome architecture. Created with BioRender.com.

Next, we explore the effect of NIPBL depletion on specification towards the three germ layers. We convert EpiSCs into neuroectoderm (NE), mesoderm (MES), and definitive endoderm (DE), using lineage-directed differentiation protocols (**Fig. 5G-I**). Robust activation of lineage-specific marker genes in all three germ layers (**Supp Fig. 10**) validates our differentiation protocols. Bulk RNA-seq analysis reveals that NIPBL depletion results in incomplete gene activation during the formation of the three germ layers, particularly for DE (**Fig. 5J-L** and **Supp Fig. 11**). DE exhibits the lowest correlation (R=0.86, versus 0.93 for MES and 0.95 for NE) in gene activation and inactivation profiles between dTAG- and DMSO-treated cells.

Although NIPBL depletion has a more modest effect on MES differentiation than DE at the transcriptome level, several important lineage specification genes (such as *Meox1*, *Mesp1*, *Bmp4*, and *Tbx6*) are insufficiently activated when NIPBL is depleted (**Supp Fig. 11B and 11E**). scRNA-seq data and immunocytochemistry results further show decreased proportion of TBX6-positive mesodermal cells in the differentiated population (**Supp Fig. 11J and 12**).

The significant downregulation of definitive endoderm marker genes, including *Sox17*, *Cxcr4*, *Gata6*, and *Gata4* (**Supp Fig. 11C and 11G**), prompts us to examine whether NIPBL depletion impairs definitive endoderm formation. scRNA-seq analysis of the cell population obtained by the DE differentiation protocol, reveals that the proportion of *Sox17*- and *Gata6*-positive definitive endoderm cells is markedly reduced upon NIPBL depletion (**Fig. 5M-O** and **Supp Fig. 13A-B**). This result is further corroborated by SOX17 immunofluorescence (**Supp Fig. 11K**). Moreover, NIPBL depletion results in distinct primitive streak-like cell populations (clusters 1, 4, and 5 in **Fig. 5M**) that, contrary to control cells (clusters 0 and 6), are characterized by *Foxa2* and *Lhx1* expression but show obvious failure to upregulate *Otx2* and *Cer1* (**Fig. 5M-O** and **Supp Fig. 13C**). The Primitive Streak (PS) is the precursor of the definitive endoderm, and key markers including *Foxa2*, *Otx2*, *Lhx1*, and *Cer1* are robustly activated within the PS^70–74^. To further confirm the existence of these abnormal Primitive Streak-like cells after NIPBL depletion, we perform a partial definitive endoderm differentiation protocol and collect PS–stage samples to analyze *Otx2* and *Foxa2* expression (**Fig. 5I**). qPCR results reveal significantly reduced *Otx2* and *Foxa2* expression in PS cells when NIPBL is depleted (**Fig. 5P**). These results indicate that NIPBL depletion results in abnormal PS-like cells and consequently compromises definitive endoderm formation.

Collectively, our findings reveal that NIPBL function is needed for activation of several genes important for lineage specification and cell differentiation.

### NIPBL mediates spatial proximity between distal enhancer and promoter to enable full activation of *Pou3f1* in EpiLCs

To explore the impact of NIPBL depletion on enhancer-promoter communication during gene activation, we establish cell lines carrying the MS2/MCP imaging system to detect the nascent transcription kinetics of *Pou3f1* during EpiLC differentiation. As *Pou3f1* is activated during the ESC to EpiLC transition, its transcriptional bursting increases (**Supp Fig. 14A-B**). Very early on (8 h), a small proportion (20%) of *Pou3f1* promoters switch on to a state permissive for bursting, exhibiting a relatively low bursting frequency (∼0.4 bursts per hour). As the transition to EpiLC progresses, the majority of *Pou3f1* promoters switch to bursting-permissive state, reaching ∼74% and ∼93% at 24 h and 48 h, respectively. The bursting frequency is also dramatically increased, during this time, reaching 3.0 and 3.3 bursts per hour at 24 h and 48 h time points, respectively (**Supp Fig. 14B**). During these significant changes in burst frequency and on-population (population of alleles in the permissive state), the burst amplitude and duration show only a slight change. The real-time nascent transcription dynamics are consistent with qPCR results showing time-dependent upregulation of *Pou3f1* (**Supp Fig. 14C**). These observations provide key insights into the dynamics of gene activation orchestrated through progressively enhanced enhancer–promoter communication during cell state transitions.

H3K27ac ChIP-seq data^75^ show several enhancers surrounding the *Pou3f1* locus that become progressively more active during *Pou3f1* activation. We focus on a distal enhancer located ∼200 kb upstream of *Pou3f1* (**Fig. 6A**), reasoning that its regulatory function is likely to depend on action-at-a-distance mechanisms. We homozygously delete this distal enhancer using clustered regularly repeating interspaced short palindromic repeats (CRISPR)/Cas9 in mESCs and then convert these cells into EpiLCs. Deletion of the distal enhancer significantly impairs *Pou3f1* activation and reduces *Pou3f1* burst probability during the EpiLCs transition (**Fig. 6B-D**), validating this distal region as a bona fide *Pou3f1* enhancer. Interestingly, although this distal 200 kb enhancer is needed for full activation at both 24 h and 48 h of the EpiLC transition, *Pou3f1* is not sensitive to NIPBL depletion at 24 h but becomes sensitive at 48 h, when NIPBL depletion results in a comparable transcriptional attenuation to deletion of the 200 kb enhancer (**Fig. 6E–F**). These results suggest that NIPBL might be critical for the interaction between the 200 kb distal enhancer and the *Pou3f1* promoter, in a time-dependent manner during the EpiLC formation.

To investigate the underlying changes in the spatial relations between *Pou3f1* promoter and the 200 kb distal enhancer, we perform multi-color super-resolution imaging with OligoPAINT FISH probes^76^ (**Fig. 6A and G**). Nanometer-scale 3D distance measurements reveal that proximity between *Pou3f1* promoter and distal enhancer increases as *Pou3f1* gets activated, with the median distance decreasing from 395 nm in mESCs to 301 nm and 281 nm in 24 h and 48 h EpiLCs, respectively (**Fig. 6H**). Consistent with the bursting results, NIPBL depletion resulted in a shift of the overall distribution toward larger distances in 48 h EpiLCs, but not in 24 h EpiLCs, with an increase of 117 nm in the median distance (**Fig. 6I-J**). These results further demonstrate that NIPBL mediates spatial proximity between the distal enhancer and promoter to enable full activation of *Pou3f1* in EpiLCs.

### NIPBL facilitates timely and precise gene activation by physically bringing newly activated enhancers and promoters into proximity

The formation of DE from EpiSCs is accompanied by wide-spread changes in the epigenetic landscape and chromatin accessibility, as differentiation signals lead to activation of new enhancers and inactivation of old ones^49^. We perform single-cell multi-omics (scMultiome) analysis on differentiated DE cells, to examine chromatin accessibility across different cell sub-populations and to investigate any effects of NIPBL depletion on epigenetic landscape changes. To pinpoint likely cis-elements important for the EpiSC to DE transition, we use the Peak2GeneLinkage function in ArchR to identify all peaks associated with activated or deactivated genes (**Supp Fig. 15A**). NIPBL depletion slightly changes chromatin accessibility of these genes in DE cells (**Supp Fig. 15B**). Since the major transcriptome changes upon NIPBL depletion occurred in PS-like cell sub-populations, we focus on differentially expressed genes (DEGs) comparing DMSO- and dTAG-treated PS-like clusters. We then overlap these DEGs with the sets of activated and deactivated genes (**Supp Fig. 15C**) and compare chromatin accessibility for the overlapping genes. Our analysis shows that there are no obvious changes in chromatin accessibility of likely *cis*-elements caused by NIPBL depletion in PS-like cells (**Supp Fig. 15B**).

Although chromatin accessibility is not affected by NIPBL depletion, genes like *Otx2* and *Foxa2* are incompletely activated during DE formation. We reason that this inability to convert epigenetic changes into transcriptional outputs might reflect disruption in distal enhancer-promoter communication. Our Hi-C and virtual 4C analysis reveals that NIPBL depletion reduces the contact frequency between the promoters and putative distal enhancers of *Otx2* and *Foxa2* (**Fig. 7A-B**). Examination of published bulk ATAC-seq data reveals increased accessibility of these putative distal enhancers of *Otx2* and *Foxa2* in DE cells compared with EpiSCs (**Fig. 7C-D**). When we delete the putative enhancers, we observe reduced *Otx2* and *Foxa2* expression and downregulation of *Sox17* during DE formation, suggesting they are indeed bona fide enhancers (**Fig. 7E-F**). The distal *Otx2* enhancer in particular appears to be specific for DE formation: although *Otx2* is already activated to a lesser extent in EpiLCs and H3K27ac levels at this distal enhancer increase, deletion of this enhancer does not affect activation of *Otx2* in EpiLCs or other EpiLC genes like *Fgf5* and *Dnmt3b* (**Supp Fig. 16**). To further understand the effect of NIPBL depletion on enhancer-promoter interactions, we use OligoPAINT FISH to measure the 3D distances between the distal enhancers and promoters of *Otx2* and *Foxa2*, specifically in Primitive Streak cells, the intermediate state when these genes are activated during the EpiSC-to-DE transition. NIPBL depletion results in an overall shift of the distance distribution toward larger values, with an increase of 55 nm and 110 nm in the median distance for *Otx2* and *Foxa2*, respectively (**Fig. 7G-H**). Overall, these results indicate that NIPBL modulates promoter-enhancer proximity and interactions to ensure full activation of *Otx2* and *Foxa2* in Primitive Streak cells, thereby guaranteeing efficient definitive endoderm specification.

Consistent with the genome-wide chromatin accessibility analysis, browser tracks with peak-to-gene links at the *Otx2* and *Foxa2* loci show that NIPBL depletion do not markedly affect the chromatin accessibility of their distal enhancers or promoters in PS-like cell sub-populations (**Fig. 7I-J**). Collectively, these results suggest that NIPBL facilitates timely and precise gene activation by physically bringing enhancers and promoters into proximity, thereby enabling cells to timely respond to developmental cues that reshape the epigenetic landscape and translate these changes into transcriptional outputs.

### A secondary, context-dependent mechanism, links NIPBL function to maintaining long-term repression of lineage-inappropriate genes

Beyond the incomplete activation of lineage-specific genes, we discover that long-term NIPBL depletion during the conversion of naïve pluripotent ESCs to EpiSCs leads to aberrant activation of a substantial set of genes that normally remain silent at this developmental stage (**Fig. 8A and Supp Fig. 17**). Gene Ontology (GO) analysis reveals that these upregulated genes are significantly enriched in biological processes related to later developmental stages, including pattern specification process, urogenital system development, and morphogenesis of a branching structure (**Fig. 8B**). Notably, genes across multiple *Hox* clusters exhibited aberrant upregulation following NIPBL depletion (**Fig. 8C**). To determine whether this aberrant upregulation of numerous genes is specific to NIPBL-dependent repression mechanisms in the final EpiSC state, or some intermediate time point during the transition, we deplete NIPBL acutely in EpiSCs or at an intermediate point when ESCs have transitioned to EpiLCs (**Supp Fig. 18A-B**). Bulk RNA-seq analysis shows that neither NIPBL depletion in EpiSCs alone nor depletion during the EpiLC-to-EpiSC transition results in a comparable number of upregulated genes to the prolonged depletion from ESCs to EpiSCs (**Supp Fig. 18A-C**). These results indicate that the aberrant upregulation of numerous lineage inappropriate genes might be a cumulative effect of the prolonged NIPBL depletion during the sequential transitions from ESC-to-EpiLC-to-EpiSC, mechanistically distinct from the NIPBL roles in enhancer-promoter communication during activation of lineage specification genes.

During epiblast differentiation, the *de novo* DNA methyltransferases DNMT3A and DNMT3B are upregulated and contribute to restraining the premature activation of lineage-specifying genes. Our bulk RNA-seq results show that NIPBL depletion results incomplete activation of *Dnmt3b* (|Fold change| ≥ 2, p-value < 0.01) (**Supp Fig. 9C**). However, the aberrant upregulation of lineage inappropriate genes upon prolonged NIPBL depletion does not appear to be a secondary effect related to reduced *Dnmt3b* expression. Inducible overexpression of DNMT3B in EpiLCs during the transition to EpiSCs using the Tet-on system does not rescue the aberrant gene upregulation (**Fig. 8D-F** and **Supp Fig. 19**).

In addition, we observe reduced contact frequencies of cohesin-mediated loops between the *HoxA* and *HoxD* cluster loci and cohesin/CTCF binding sites (**Fig. 8G and Supp Fig. 20**), with similar decreases also observed at the *HoxB* cluster (data not shown).

Taken together, these results suggest an association between disrupted cohesin-mediated chromatin interactions and the derepression of lineage-specifying genes upon NIPBL depletion during epiblast differentiation.

## Discussion

Early embryonic development highly depends on the rapid and precise regulation of gene-expression programs to specify lineage identity and guide cell-fate commitment. Genome architecture is proposed to scaffold hierarchical regulatory landscapes, providing a framework for communication between regulatory elements and shaping spatial and temporal control of gene expression in response to developmental cues. Here, we demonstrate that NIPBL, and thereby cohesin-mediated genome folding, is indispensable for orchestrating gene reprogramming during the transition from naïve pluripotency to definitive endoderm. Mechanistically, integrated Hi-C and single-gene imaging analyses illustrate that NIPBL directly regulates enhancer-promoter interactions by physically bringing them in proximity (**Fig. 9**).

By leveraging an advanced *in vitro* differentiation system, we are able to interrogate the progressive changes in 3D genome architecture across early embryogenesis, spanning the transition from pre-implantation epiblast to post-implantation gastrulation—a critical window of mammalian development. Our genome-wide analysis demonstrates that chromatin loops are different among naïve pluripotency to primed pluripotency and finally to definitive endoderm. This reflects the distinct transcriptional programs among the three cell types and underscores the important regulatory role of chromatin loops^77^. Additionally, changes in compartment interactions indicate shifts in the spatial segregation of active and inactive chromatin into their respective compartments^78^. In contrast, TAD structures remain largely conserved across the three cell types, consistent with previous observations^79–81^.

*In vivo*, the transitions in 3D genome architecture across these three cell types occur rapidly during early mouse embryonic development, approximately from embryonic day (E) 3.5 to E7.5, and therefore this reorganization requires tight and precise modulation. Cohesin has been established as a key architectural regulator through its loop-extrusion activity; however, its multiple essential functions in cell cycle progression and sister chromatid segregation during mitosis makes it a less suitable target for probing how genome architecture influences cell-fate maintenance and transitions. Therefore, we focused on NIPBL, a protein important for the loop extrusion activity of cohesin, whose reduced dosage—unlike cohesin—does not disrupt mitotic progression, as reported previously^44^ and confirmed by our own observations. In addition, our Hi-C analyses show that NIPBL depletion results in architectural alterations that mirror those caused by cohesin depletion, including increased compartmentalization, reduced TAD contacts, and diminished chromatin loop contact frequencies, although the overall effects are milder than those observed after direct cohesin removal^17,33^. Furthermore, these NIPBL-dependent architectural changes are remarkably consistent across the three cell types examined, aligning with the conserved effects reported for NIPBL depletion in mouse liver and mononuclear neutrophil progenitors^82,83^. Altogether, these results support the conclusion that NIPBL functions as a conserved regulator of genome architecture by facilitating cohesin-mediated loop extrusion across distinct cellular states.

By performing scRNA-seq and single-cell multi-omics, we resolve cellular heterogeneity and comprehensively delineate the phenotypic consequences of NIPBL depletion on cell-state maintenance and lineage transitions during early embryonic development. A recently published study reports that cell-fate transitions, such as astrocyte differentiation or gastruloid formation, remain largely robust despite NIPBL depletion, based on bulk RNA-seq and bulk ATAC-seq analyses ^84^. However, these bulk-derived conclusions seem difficult to reconcile with the reduced mesoderm cell numbers, the altered mesodermal subpopulation proportions in *Nipbl*-haploinsufficient mouse embryos, and the severe developmental abnormalities observed in both mouse and human NIPBL mutants^37–39,85^. In addition, subtle transcriptional perturbations caused by NIPBL haploinsufficiency in both mice and human cells have been shown to result in severe developmental defects and disease^39,86^. Together, these findings highlight the importance of considering cellular heterogeneity and the regulatory role of NIPBL in controlling key cell-fate genes.

In our study, we find that NIPBL depletion reduces the efficiency of definitive endoderm formation, which is caused by cells undergoing altered differentiation trajectories. These altered trajectories result from the incomplete activation of key lineage-specifying genes, such as *Otx2*^71,72^ and *Foxa2*^87–89^, leading cells to transit through distinct primitive streak–like intermediate states. Consistent with the mesodermal defects reported in *Nipbl*-haploinsufficient mouse embryos, we observe a markedly reduced proportion of mesodermal cells after NIPBL depletion. By contrast, the transition toward the neuroectodermal lineage remains largely robust. The pronounced differences among the three germ layers derived from the same starting cell population and at the same differentiation time point may reflect the distinct trajectory histories experienced by the cells before reaching these fates. Furthermore, this interpretation is further supported by the progressively worsening defects observed during the successive ESC-to-EpiSC transitions.

Mechanistically, by integrating microscopy-based methods with chromosome conformation capture, two complementary technologies central to 3D genome research^77^, we demonstrate that NIPBL is essential for cohesin-mediated enhancer-promoter communication by facilitating their physical proximity through loop extrusion, thereby ensuring proper gene activation. Although cohesin has long been proposed to regulate transcription through its architectural role, particularly by organizing enhancer–promoter contacts, several studies have reported that acute cohesin depletion results in only modest transcriptional changes, leading to debate over its regulatory function^11,33^. Our results help reconcile this discrepancy by providing convergent evidence from both imaging and high-throughput sequencing that cohesin is indeed required for enhancer–promoter interactions and plays a specific regulatory role in gene expression during cell-fate maintenance and transitions. Building on these findings, previous studies have demonstrated that NIPBL regulates transcription through interactions with mediator, BRD4, and various transcription factors^90–93^, and we speculate that such interactions could facilitate the precise deployment of cohesin at regulatory elements to modulate enhancer–promoter communication. This hypothesis will require further experimental testing.

Furthermore, our findings suggest that NIPBL plays a critical role in stabilizing enhancer networks required for de novo gene activation. For example, activation of *Pou3f1* relies on multiple distal enhancers, and its expression fails in a substantial fraction of cells or alleles when NIPBL is depleted or when the most distal enhancers are removed. This behavior points to highly non-additive interactions among enhancers, raising the possibility that perturbing NIPBL disrupts the cooperative architecture of these elements, causing the entire enhancer network to collapse at the single-cell or single-allele level. In line with this model, our multi-omics analyses indicate that differentiating cells still receive inductive cues and initiate upstream signaling cascades, but these cues fail to be transmitted to the promoter when NIPBL is absent. Although new *cis*-regulatory elements become accessible, as revealed by scATAC-seq, their activation does not propagate to productive transcriptional output, suggesting that NIPBL is essential for relaying lineage-specific signals from enhancers to promoters during gene activation.

In addition to facilitating gene activation, cohesin-associated CTCF sites have also been reported to function as insulators that may repress the expression of lineage-specifying developmental genes, such as Hox cluster genes^94,95^. In our study, we find that a prominent feature of long-term NIPBL depletion is the broad derepression of genes in EpiSCs. Given the critical role of *de novo* DNA methylation in stabilizing gene repression in the primed pluripotent state^96,97^, we initially attributed this derepression to the marked downregulation of the DNA methyltransferase *Dnmt3b*. However, restoring DNMT3B expression in NIPBL-depleted EpiSCs failed to rescue the abnormal derepression, suggesting that additional architectural mechanisms are likely contributing to this phenotype. Beyond DNA methylation, Polycomb complexes are also implicated in maintaining the repression of key developmental genes, in part through the formation of Polycomb domains that are reinforced by PRC1-mediated long-range chromatin interactions^98,99^. Consistent with previous studies demonstrating that cohesin counteracts Polycomb-mediated chromatin domain interactions^100–102^, depletion of NIPBL in EpiSCs led to strengthened long-range Polycomb loops at *Hox* cluster loci (not shown). However, we observed a decrease in cohesin-mediated loop contacts near *Hox* cluster loci upon NIPBL depletion. These observations support the possibility that Nipbl contributes to proper gene regulation by facilitating cohesin-mediated genome folding. This finding warrants further investigation into how cohesin-driven loop extrusion participates in gene repression and the mechanisms underlying this regulation.

In summary, our study provides a more comprehensive insight into the role of cohesin-mediated 3D genome architecture in helping establish and maintain gene expression programs crucial for specifying cell fates and identities during early embryonic development. This work provides a framework linking cohesin-mediated loop extrusion to developmental potential and sets the stage for future studies dissecting how genome conformation contributes to mammalian embryonic development.

## Data availability

Raw bulk RNA-seq, Hi-C, and single-cell RNA-seq and single-cell Multiome data generated in this study have been deposited in the Sequence Read Archive (SRA) under the accession number (PRJNA1390683). Published RAD21 and H3K27ac ChIP-seq datasets in mESCs (GSE183828), H3K27ac ChIP-seq datasets in EpiLCs (GSE117896), bulk ATAC-seq datasets in EpiSCs and definitive endoderm cells (GSE189869), and CTCF and SMC3 ChIP-seq datasets in E6.5 epiblast (GSE200323) were obtained from the Gene Expression Omnibus (GEO). Raw data were reanalyzed as described in the Methods.

## Code availability

All analyses were performed with standard, publicly available tools as described in Methods.

## Acknowledgements

We thank Luke Lavis (HHMI/Janelia) for dye-labeling reagents, and Ronan Chaligné for advice on single-cell multi-omics experiments. Single-cell multi-omics experiments were performed in the MSK Single-cell Analytics Innovation Lab (SAIL). We thank Christina Leslie and Effie Apostolou, as well as the members of the Pertsinidis lab for helpful discussions. This work is supported by the National Institute of General Medical Sciences of NIH (grants R01GM135545, R21GM134342, and R01GM144508 (AP)), the Tri-Institutional Stem Cell Initiative supported by The Starr Foundation (AP), the SKI Basic Research Innovation Award Initiative (BRIA) and the Dewitt Wallace Basic Science Award Fund (LC and AP), and the National Cancer Institute (grant P30 CA008748).

## Competing interests

The authors declare no competing interests.

## Author Contributions

Z.N. and A.P. conceived the project. X.M. performed all bioinformatic analyses. Z.N., S.D., L.C., R.G., T.V. and A.P. designed the research, performed the experiments, analyzed the data and interpreted the results. T.V. and A.P. supervised the project. Z.N. and A.P. wrote the manuscript with input from all coauthors.

## Methods

### Cell lines

Mouse embryonic stem cell lines were derived from Bruce 4 mESCs (Millipore Sigma CMTI-2; murine strain C57/BL6J, male) and a strain C57BL/6J (B6) mouse blastocyst (ATCC, SCRC-1002; male). Mouse embryonic fibroblasts (MEFs) were isolated from the stroma of C57BL/6 mice (ATCC, SCRC-1008).

### Cell culture condition and differentiation

All cells were cultured at 37 °C in a humidified atmosphere with 5% (v/v) CO_2_. mESCs were maintained on 0.1% gelatin-coated dishes with standard serum/2i medium containing DMEM (Gibco, 10313039), 15% fetal bovine serum (R&D Systems, S10250), 1000 U/mL LIF (Millipore, ESG1107), 2 mM L-alanyl-L-glutamine (Gibco, 35050079), 1× MEM nonessential amino acids (Gibco, 11140076), 0.1 mM 2-Mercaptoethanol (Gibco, 21985023), 3 μM CHIR99021 (Millipore, 361559), 1 μM PD0325901 (Axon Medchem, 1408) and 100 U/mL Penicillin-Streptomycin (Gibco, 15140122).

EpiLC, EpiSC, and three germ layer differentiation was induced as previously described^49,50,103^. Briefly, mESC colonies were dissociated with 0.025% trypsin-EDTA (Sigma, SM-2004-C) and plated at a density of 1.8 ×10^4^ cells/cm2 on Fibronectin (Sigma, FC010) coated dishes in N2B27-based medium supplemented with 1% KnockOut Serum Replacement (KSR) (Gibco, 10828010), 12.5 ng/mL bFGF (Thermo Fisher Scientific, PHG0367), and 20 ng/mL activin A (Peprotech, 120-14P).

For EpiSC differentiation, 48h EpiLCs were lifted with Accutase (Gibco, A1110501) and seeded at a density of ∼1×10^5^ cells/cm^2^ on mitomycin C-treated MEFs in N2B27-based medium supplemented with 175 nM NVP-TNKS656 (Selleck Chemicals, S7238), 20 ng/mL activin A, and 12.5 ng/mL bFGF.

For definitive endoderm differentiation, EpiSC colonies were dissociated with Collagenase IV (Gibco, 17104109) and Accutase, and seeded onto rhlaminin-521-coated (Gibco, A29249) dished in plating medium composed of chemically defined medium (CDM) supplemented with 0.7 ng/μL insulin, 1% KSR, 175 nM NVP-TNKS656, 12.5 ng/mL bFGF, 20 ng/mL activin A, and 2 μM Thiazovivin (Sigma, SML1045). Plating medium was replaced 6 h after seeding with CDM supplemented with 0.7 ng/μL insulin, 40 ng/mL activin A and 3 μM CHIR99201 for 16 h, followed by CDM supplemented with 0.7 ng/μL insulin, 100 nM LDN-193189 (Stemgent, 04-0074) and 100 ng/mL activin A for an additional 24 h.

For mesoderm differentiation, EpiSC colonies were dissociated same as above, and seeded onto rhlaminin-521-coated dished in plating medium. Plating medium is composed of CDM plus 0.7 ng/μL insulin medium supplemented with 1% KSR, 175 nM NVP-TNKS656, 12.5 ng/mL bFGF, 20 ng/mL activin A, 5 µM Emricasan (Selleckchem, S7775), and 50 nM Chroman 1 (MedChem Express, HY-15392). Plating medium was replaced 6 h after seeding with CDM plus 0.7 ng/μL insulin medium supplemented with 25 ng/mL bFGF and 3 μM CHIR99201 (differentiation medium). After 16 h, the medium was replaced with fresh differentiation medium for an additional 24 h.

For neuroectoderm differentiation, EpiSC colonies were dissociated same as above, and seeded onto rhlaminin-521-coated dishes in N2B27-based medium (B27 minus vitamin A; Gibco, 12587010) supplemented with10 μM SB431542 (Tocris, 1614) and 100 nM LDN-193189 (differentiation medium). After 22 h, the medium was replaced with fresh differentiation medium for an additional 24 h.

### Degron cell line generation and targeted protein degradation

*Nipbl*-dTAG-Halo and *Rad21*-dTAG-Halo cell lines were generated as described previously^61^. Specifically, an HA-Halo-FKBPF36V tag was inserted at the N terminus of NIPBL immediately after the start codon, and at the C terminus of RAD21 immediately before the stop codon. dTAG-13 (Sigma, SML2601) was dissolved in DMSO (Thermo Fisher Scientific, D12345) to prepare a 2 mM stock solution, and cells were treated with 400 nM dTAG-13 for protein degradation.

### Hi-C

Hi-C and library preparation was performed using the Arima-HiC kit (Arima, A510008) and the Arima Library Prep Kit (Arima, A303011) according to the manufacturer’s instructions. Prior to Hi-C experiments, mESCs and EpiSCs were treated with dTAG for 4 h to induce NIPBL depletion. For definitive endoderm, dTAG treatment was initiated at the EpiSC stage for 4 h and maintained throughout subsequent differentiation. mESCs and EpiSCs were detached and dissociated into single cells as described above, while definitive endoderm cells were dissociated with Accutase as described previously^49^. Dead cells were removed using the Miltenyi Dead Cell Removal Kit (Miltenyi Biotec, 130090101), and cell viability was assessed using the automated cell counter. Hi-C was performed using ∼2×10⁶ cells with >90% viability per replicate. Libraries were sequenced on an Illumina NovaSeq 6000 platform using paired-end 150 bp reads.

### MS2/MCP cell line generation

The *Klf4*/*Tbx3*/*Sox2*-MS2/MCP–mNeonGreen ESC lines were established as described previously^61,104^. Specifically, 24×MS2 stem-loop cassettes were inserted into the 3′ UTRs of both alleles of *Klf4* and *Tbx3*, and into one allele of *Sox2*. The knock-in cells constitutively expressed MCP–mNeonGreen through transposase-mediated genomic integration by co-transfecting pPB-LR5-CAG-MCP-mNeonGreen-IRES-Neo and pCMV-hyPBase plasmids using Lipofectamine 2000 (Invitrogen, 11668019).

*Pou3f1*-MS2/MCP–mNeonGreen ESC lines were generated following the same procedure as for the *Klf4*/*Tbx3*/*Sox2*-MS2/MCP–mNeonGreen lines. Specifically, to perform selection in mESCs, a PGK promoter–driven Hygromycin resistance (HygR) cassette was designed downstream of the 24×MS2 cassette. Transfected cells were selected with 100 μg/mL Hygromycin until all untransfected cells were eliminated. After selection, the HygR cassette flanked by loxP sites in the same orientation was excised by transient transfection with Cre expression vectors using Lipofectamine 2000. Homozygous clones were identified by PCR genotyping. Fluorescence-activated cell sorting (FACS) was performed to remove MCP–mNeonGreen–negative cells.

### LacO/LacI-SNAP and TetO/TetR-Halo NIPBL degron cell line generation

These cell lines were established following the same procedure as the *Pou3f1*-MS2/MCP–mNeonGreen lines, and NIPBL degron integration was then performed as described above. For the *Sox2* locus, a 48×TetO array was integrated at chr3:34,755,422, located ∼100 kb downstream of the *Sox2* gene, where the downstream enhancers are located, and a 90×LacO array was inserted at chr3:34,646,417 near the promoter. For the *Klf4* locus, a 96×TetO array was integrated at chr4:55,483,317, located ∼55 kb downstream of the *Klf4* gene, where the downstream enhancers are located, and a 90×LacO array was inserted at chr4:55,535,211 near the promoter. To perform selection, PGK promoter–driven resistance cassettes were designed downstream of the arrays: a Neo cassette flanked by Frt sites for *Sox2*-enhancer 48×TetO targeting; a Hygromycin (HygR) cassette flanked by Rox sites for *Sox2*-promoter 90×LacO targeting; a Neo cassette flanked by LoxP sites for *Klf4*-enhancer 96×TetO targeting; and a Puromycin (Puro) cassette flanked by Frt sites for *Klf4*-promoter 90×LacO targeting. Cells were selected with 400 µg/mL G418 (Sigma, G8168), 1 µg/mL puromycin, and 100 µg/mL Hygromycin. To excise the selection cassettes, pFlipase, pDre, and pCre expression vectors were transiently transfected using Lipofectamine 2000. PCR genotyping was performed to identify knock-in clones.

HA–CLIP–FKBPF36V tags were inserted at the N terminus of NIPBL, immediately after the start codon, in the *Klf4*-promoter-90×LacO–enhancer-96×TetO and *Sox2*-promoter-90×LacO–enhancer-48×TetO cell lines, respectively.

To isolate NIPBL-CLIP-, LacI-SNAP-, and TetR-Halo-positive cells by FACS, cells were stained with 0.6 μM JF549-CLIP, 0.3 μM JFX650-SNAP, and 0.3 μM JF646-Halo dyes (Janelia Fluor®) in culture medium prior to sorting.

### Targeting vector construction

The donor plasmids used for endogenous targeting were designed as described previously^61^ and synthesized by GenScript. Because the repeat arrays could not be synthesized, they were inserted into the vectors by restriction enzyme digestion and ligation. The 24×MS2 cassette was excised from the pCR4-24×MS2SL-stable plasmid (Addgene #31865), the 48×TetO cassette from the 48mer EFS-blaR EFS-hsvTK plasmid^105^, and the 90×LacO cassette from the pLAU43 plasmid^106^.

### Live cell imaging

Mouse ESCs were plated in standard serum/2i medium onto 8-chamber coverglass (Nunc™ Lab-Tek™ II Chambered Coverglass, 155409) pre-coated with 5 µg/mL laminin-511 (BioLamina, LN511). Prior to imaging, cells were treated with DMSO or dTAG-13 for 4 h to degrade target proteins.

For EpiLC imaging, mouse ESCs were dissociated and replated in EpiLC differentiation medium onto laminin-511-coated chamber coverglass to continue differentiation.

Real-time tracking of nascent transcription using the MS2/MCP system was performed as previously described^61^. Briefly, live-cell imaging was performed on a Zeiss AxioObserver wide-field epifluorescence microscope equipped with a 63×/1.4 NA oil-immersion objective in a humidified chamber at 37 °C and 5% CO₂. The samples were imaged at 45- or 60-second intervals using 260-nm z-steps. Images were acquired in 16-bit mode at a resolution of 1024 × 1024 pixels, with a pixel size of 206 nm.

For live-cell measurement of *Klf4* and *Sox2* enhancer–promoter (E–P) distances in mouse ESCs, imaging was performed on a Leica TCS SP8 confocal microscope equipped with a 100×/1.44 NA oil-immersion objective (Leica HC PL APO) in a humidified chamber maintained at 37 °C with 5% CO₂. Prior to imaging, cells were labeled in culture medium with 0.3 μM JF554-Halo and 0.3 μM JFX650-SNAP dyes. Dual-color images were simultaneously acquired using two SPCM-AQR (PerkinElmer) avalanche photodiode (APD) detectors with band-pass filters (Chroma ET595/50 m and ET690/50 m) for TetR-Halo and LacI-SNAP emissions, respectively. Images were collected in 12-bit mode at a resolution of 128 × 128 pixels, with a pixel size of 30.5 nm and z-steps of 100 nm.

### Mouse ESCs synchronization and imaging

*Nipbl*-dTAG-Halo and *Rad21*-dTAG-Halo mESCs were seeded in standard serum/2i medium onto 8-chamber coverglass pre-coated with laminin-511 and cultured overnight. Cells were arrested in mitosis by treatment with 200 ng/mL Nocodazole (Sigma, M1404) for 6 h, and subsequently released by washing 6 times with fresh culture medium to remove Nocodazole. To degrade target proteins, dTAG-13 was added 4 h before release.

For DNA labeling, cells were incubated with 500 nM SiR-DNA probe (Spirochrome, SC007) at 37 °C for 1 h prior to release, and unbound probes were removed by washing with fresh medium during the release step. DNA tracking was performed as described in the live-cell imaging section. Images were acquired at 2-min intervals with 900-nm z-steps.

### gRNA design and validation

The guide RNAs (gRNAs) were designed using the CRISPR Design Tool. Oligonucleotide pairs were annealed and cloned into the BbsI-digested eSpCas9 plasmid (Addgene, 71814). The efficiency of each gRNA was validated by Surveyor assay as described previously^107^. All gRNAs used in this study are listed in Table S3.

### Enhancer deletion

For enhancer deletions, two gRNAs targeting the left and right boundary of their respective H3K27ac ChIP-Seq (data from Pengyi Yang et al., 2019) and ATAC-seq peaks (data from Daniel Medina-Cano et al., 2022) were designed using the CRISPR Design Tool. Cloning and validation of gRNA efficiency were performed as described above. Approximately 2×10^5^ mouse ESCs were plated in one well of a 6-well plate and transiently transfected the next day with 1 µg of each gRNA plasmid using Lipofectamine 2000 following the manufacturer’s protocol. Transfected cells were selected with 1 µg/mL puromycin for 24 hours, and puromycin-resistant colonies were picked and screened by PCR genotyping to identify homologous clones.

### OligoPAINT DNA FISH probe design and DNA FISH

OligoPAINT DNA FISH probes were designed as described previously^61^. In this study, the primary DNA FISH probes were designed against the following genomic loci: *Pou3f1* promoter (chr4:124654820-124658234), *Pou3f1* distal enhancer (chr4:124441978-124445392), *Otx2* promoter (chr14:48667620-48671000), *Otx2* distal enhancer (chr14:48704414-48709864), Foxa2 promoter (chr2:148045938-148049911), *Foxa2* distal enhancer (chr2:148097125-148101098). In addition, four supporting probe pools were designed flanking the target loci to facilitate the identification of each gene locus. The coordinates of the supporting probes were as follows: *Pou3f1* probe1 (chr4:124593137-124606796), *Pou3f1* probe2 (chr4:124703216-124716875), *Pou3f1* probe3 (chr4:124393210-124406869), *Pou3f1* probe4 (chr4:124513196-124526855), *Otx2* probe1 (chr14:48579102-48590003), *Otx2* probe2 (chr14:48619233-48630134), *Otx2* probe3 (chr14:48799532-48810433), *Otx2* probe4 (chr14:48744671-48755572), *Foxa2* probe1 (chr2:147956075-147964021), *Foxa2* probe2 (chr2:148001123-148009069), *Foxa2* probe3 (chr2:148186031-148193977), *Foxa2* probe4 (chr2:148146055-148154001). Each primary probe set was composed of 55 individual oligonucleotide probes, designed based on the mm10 reference genome. The probe pools were synthesized by Integrated DNA Technologies (IDT).

EpiLC DNA FISH samples were prepared in the same way as live-cell imaging samples. For primitive streak sample preparation, EpiSC colonies were dissociated with Collagenase IV and Accutase, and seeded onto rhLaminin-521–coated 8-chamber coverglasses in plating medium. The plating medium was replaced 6 hours after seeding with CDM supplemented with 0.7 ng/μL insulin, 40 ng/mL activin A and 3 μM CHIR99201 for 16 h.

DNA FISH was performed as described previously^61^. The readout probe was a 51-nt oligonucleotide containing a sequence complementary to the primary probes at one end and a sequence complementary to the imaging probe at the other end. Promoter imaging probes were labeled with Cy3, enhancer imaging probes with Cy5, and supporting imaging probes with Atto 488.

DNA FISH samples were imaged as previously described^61^ using a Leica TCS SP8 confocal microscope equipped with a 100×/1.44 NA oil-immersion objective (Leica HC PL APO) at 25 °C.

### Inducible DNMT3B overexpression

The *Dnmt3b* coding sequence was amplified by PCR from EpiSC cDNA using a pair of primers containing overlap region, and the vector backbone was amplified from the pB-Halo IRES-eGFP plasmid (Addgene #164519) using a high-fidelity DNA polymerase. The CmR and eGFP cassettes were removed from the original pB-Halo IRES-eGFP vector during backbone amplification. The DNMT3B overexpression construct (pB-Halo-Dnmt3b OE vector) was assembled using the NEBuilder® HiFi DNA Assembly Master Mix (NEB, E2621S) following the manufacturer’s instructions. The integrity and accuracy of the assembled plasmid were verified by Nanopore sequencing.

Approximately 1×10^6^ homozygous Nipbl-dTAG-SNAP B6 mESCs were transfected with 6 μg of pB-Halo-*Dnmt3b* overexpression vector and 1 μg of pCMV-hyPBase plasmid using Lipofectamine 3000 (Invitrogen, L3000015) following the manufacturer’s protocol. Cells were selected with 400 µg/mL G418 (Sigma, G8168), and two single clones were picked up. After induction with 2 µg/mL doxycycline for 24 h, EpiLCs were stained with 0.3 µM JF646-Halo dye to confirm homogeneous expression of DNMT3B-Halo expression under a wide-field fluorescence microscope.

### qRT-PCR

qRT-PCR was performed as previously described^108^. Total RNA was extracted from cells that were detached as described above using TRIzol reagent (Invitrogen, 15596026) according to the manufacturer’s protocol. cDNA was synthesized using the SuperScript™ IV VILO™ Master Mix (Invitrogen, 11766050). qRT-PCR was performed using the Power SYBR™ Green PCR Master Mix (Thermo Fisher Scientific, 4367659) on an Applied Biosystems 7500 Real-Time PCR System. Primers used in the present study have been listed in Table S2.

### Single cell RNA-seq

mESCs were treated with dTAG for 6 days to induce NIPBL depletion. For mesoderm differentiation, dTAG treatment was initiated at the EpiSC stage for 4 h and maintained throughout subsequent differentiation. mESCs were dissociated using 0.025% trypsin–EDTA, whereas mesoderm cells were dissociated with Accutase, as previously described^50^. Dead cells were removed with the Miltenyi Dead Cell Removal Kit, and cell viability was assessed using an automated cell counter. The resulting cell suspensions were then filtered through a flow cytometry cell strainer (Corning, 352235). Single-cell RNA-seq libraries were prepared using the 3’ Chromium GEM-X kit (10X Genomics, v4) and sequenced in paired-end mode (PE28/88) with an average coverage of ∼20,000 reads per cell. The fastq files were processed with Cell Ranger pipeline (10x genomics cloud).

### Bulk RNA-seq

Total RNA was extracted from cells that were detached as described above using TRIzol reagent (Invitrogen, 15596026) according to the manufacturer’s protocol. RNA sequencing was performed on an Illumina NovaSeq 6000 platform using paired-end 100 bp reads.

### Single cell multi-omics

Definitive endoderm cells were prepared in the same manner as mesoderm cells for scRNA-seq. After preparation, cells were permeabilized using the DOGMA-seq DIG permeabilization protocol^109^ and subsequently processed according to the Chromium Next GEM Single Cell Multiome ATAC + Gene Expression user guide (10x Genomics, v1). Libraries were sequenced with paired end reads (PE28/88 for GEX and PE50 for ATAC) at a depth of ∼20,000 reads per cell for GEX and 25,000 reads per cell for ATAC. The fastq files were processed using Cell Ranger ARC pipeline (10x genomics, v2.0.2).

### Immunofluorescence

Immunofluorescence staining for the three germ layers was performed as previously described^49^. Primary antibodies against SOX17 (R&D Systems, AF1924), TBX6 (R&D Systems, AF4744), and SOX1 (R&D Systems, AF3369) were used in the present study, with detailed information in Table S4. Fluorescent dye-conjugated secondary antibodies (R&D Systems, NL003) were used for detection.

### Western blot

Western blot (WB) was performed as previously described^108^. In brief, cellular proteins were extracted and separated by sodium dodecyl sulfate–polyacrylamide gel electrophoresis (SDS–PAGE), followed by transfer onto a polyvinylidene difluoride (PVDF) membrane. The membrane was blocked with 5% nonfat milk, incubated with primary antibodies overnight at 4 °C, and then with HRP-conjugated secondary antibodies (Abclonal; 1:5000) for 1 h at room temperature. Protein signals were detected using the SuperSignal™ West Dura Trial kit (Thermo Fisher Scientific, 37071) according to the manufacturer’s instructions. Primary antibodies against HA (Abclonal, AE008), NIPBL (Santa Cruz, sc-374625), and DNMT3B (CST, 48488T) were used, as summarized in Table S4.

### Transcriptional bursting analysis

Analysis of transcriptional burst images was performed as previously described^61^. To avoid confounding effects from mitotic cells, only interphase cells were analyzed.

### Enhancer-promoter distance measurement

3D enhancer–promoter distances in OligoPAINT DNA FISH were measured as previously described^61^. Briefly, the xy positions of the Cy3 and Cy5 spots were extracted by least-squares fitting to 2D Gaussian functions. The z position was obtained by fitting the corresponding z-axis intensity profile, centered on the xy position, to a 1D Gaussian function with a linear background. The 3D distance between the Cy3 and Cy5 signals was subsequently calculated from their xyz coordinates.

2D enhancer–promoter distances in live-cell imaging were measured using the same procedure as for 3D distance measurements, except that the z position was not included in the calculation.

Chromatic aberrations resulting in apparent shifts between Cy3 and Cy5 images were corrected using 0.1-μm TetraSpeck beads (Invitrogen, T7279) diluted 1,000-fold in PBS to calculate the 3D offsets between the two channels.

### Gene Ontology analysis

Gene Ontology (GO) analysis was performed using the DAVID Bioinformatics Resources (https://david.ncifcrf.gov/).

### Hi-C data analysis

Hi-C contact pairs were identified using the HiC-Pro pipeline (v3.1.0) with the mm10 reference genome. The pipeline was run with default parameters to perform read alignment, filtering, deduplication, and valid pair assignment, and the resulting interaction pairs were used for contact map construction. Detailed protocols and source code are available at https://github.com/nservant/HiC-Pro.

Hi-C contact matrices were analyzed in R (v4.4.1) using the GENOVA package (v1.0.1) to investigate chromatin compartments and topologically associating domains (TADs). Compartment scores were calculated at 500 kb resolution for three cell types—ESC, EpiSC, and DE—and correlated with cell type-specific epigenetic peaks to generate signed scores distinguishing A and B compartments: ESC (GSM5571876 H3K27ac), EpiSC (SRR17074065 ATAC), and DE (SRR17074076 ATAC). TADs were identified at 10 kb resolution by computing insulation scores, and intra-TAD interactions were visualized using aggregate TAD analyses (ATA). Comparisons of TAD boundaries between samples were performed with the TADcompare R package (v1.16.0).

Chromatin loops were detected using Chromosight (v1.6.3) and Mustache (v1.2.0) at 10 kb resolution. Loops were further analyzed via aggregate peak analysis (APA) and visualized with GENOVA. Correlations of loop scores were assessed for loops overlapping cohesin-associated loops reported by Tsung-Han S. Hsieh et al., Nature Genetics, 2022. Chromatin state enrichment at loop anchors was evaluated using ChromHMM, based on segmentation states from Guifeng Wei’s publicly available mESC mm10 model (ChromHMM_mESC_mm10). Virtual 4C profiles for selected loci (*Klf4*, *Tbx3*, *Sox2*, *Otx2*, *and Foxa2*) were generated from 5 kb Hi-C matrices using Cooltools (v0.7.1).

### Single-cell RNA-seq data analysis

Single-cell RNA-seq data were processed using Seurat (v5.1.0). Low-quality cells were filtered based on gene counts (200–6000 per cell) and mitochondrial gene content (<10%). The 2,000 most highly variable genes were used for PCA-based dimensionality reduction. Cells were clustered and visualized using UMAP, with cluster annotations assigned based on known marker genes. Cell cycle status was inferred using Seurat’s CellCycleScoring function. Expression patterns of marker genes were visualized across clusters using Seurat’s VlnPlot, DotPlot, and FeaturePlot functions.

For RNA velocity analysis, spliced and unspliced counts were processed in scVelo (v0.3.3). After filtering and selecting the top 2,000 highly variable genes, RNA velocities were estimated and projected onto the UMAP embedding obtained from Seurat to visualize potential future cell states.

### Bulk RNA-seq data analysis

For RNA-seq analysis, transcript quantification was performed using StringTie (v2.2.1), and differential expression analysis was conducted with DESeq2 (v1.46.0). Genes with p < 0.01 and fold change ≥ 2 were considered differentially expressed. Volcano plots and correlation scatter plots were generated using ggplot2 (v3.4.3) in R (v4.4.1).

### Single cell multi-omics analysis

Single-cell multiome data, including scATAC-seq and matched scRNA-seq, were analyzed using ArchR (v1.0.3) with the mouse mm10 genome as reference. scATAC-seq fragments were filtered to retain cells with ≥1,000 fragments and a TSS enrichment score ≥4. scRNA-seq feature-barcode matrices were matched to scATAC-seq cells, and only cells present in both modalities were retained. The datasets were integrated into an ArchRProject, and doublets were identified and removed. Following quality control and integration, cells were clustered based on their scRNA-seq profiles. For each RNA-defined cluster, scATAC-seq data were aggregated to generate pseudo-bulk accessibility profiles. Differentially accessible regions (DARs) between clusters were identified using a Wilcoxon test, correcting for TSS enrichment and fragment count biases. Peak-to-gene correlations were calculated using ArchR’s addPeak2GeneLinks function, and chromatin accessibility and peak-to-gene links were visualized using browser tracks.

### Published ChIP-seq and bulk ATAC-seq data analysis

Raw paired-end ChIP-seq and ATAC-seq reads (FASTQ format) were processed using fastp (v0.23.4) to remove adapter sequences and low-quality bases with default parameters. Cleaned reads were aligned to the mouse reference genome (mm10) using Bowtie2 (v2.4.1). The resulting alignments were sorted and converted to BAM format using SAMtools (v1.9). PCR duplicates were identified using Picard MarkDuplicates (v2.18.29) or sambamba (v0.6.6), as indicated. For ATAC-seq samples, additional filtering was performed to remove mitochondrial reads (chrM), low-quality alignments (MAPQ <30), unmapped reads, non-primary alignments, PCR duplicates, and reads failing platform quality checks. Only properly paired reads were retained. Genome-wide signal tracks were generated from filtered BAM files using deepTools bamCoverage (v3.5.6) and output in bigWig (bw) format. Peak calling was not performed, as the analysis focused on qualitative visualization of chromatin accessibility and binding profiles. The resulting normalized signal tracks were visualized using the Integrative Genomics Viewer (IGV) (2.18.0).

### Statistics

Statistical analyses were performed using GraphPad Prism 10 (GraphPad Software) and MATLAB (MathWorks). For qPCR and other normally distributed data, comparisons between two groups were performed using unpaired two-tailed Student’s t-tests, and one-way ANOVA followed by Bonferroni’s post-hoc test was applied for multiple comparisons. P < 0.05 was considered statistically significant. All experiments were independently repeated at least three times. For imaging-based measurements, non-parametric Wilcoxon rank-sum tests were used in MATLAB. Experiments were performed with multiple independent biological replicates, as specified in the corresponding figure legends.

---

**Figure S1.**
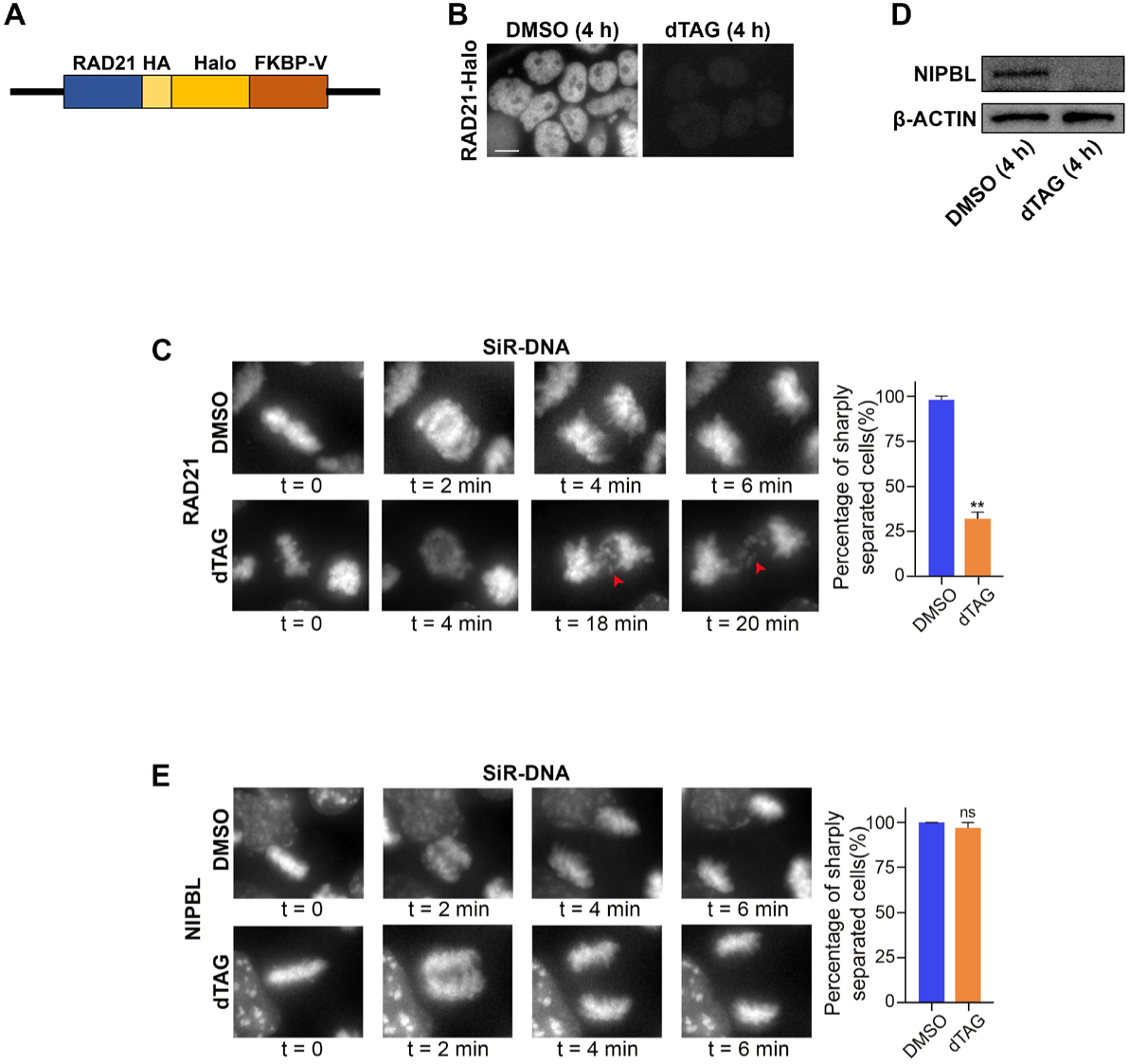
RAD21 and NIPBL depletion efficiency and effects on sister chromatid segregation. **(A)** Schematic overview of endogenous editing RAD21 for degradation using the dTAG degron system. **(B)** Representative fluorescence images showing degradation of RAD21-Halo. Scale bar = 10 µm. **(D)** Western blot analysis of NIPBL in mESCs acutely treated with DMSO or dTAG for 4 h. β-ACTIN was used as a loading control. **(C)(E)** Time-lapse images showing SiR-DNA-labeled DNA segregation during mitosis in live mESCs after acute (4 h) depletion of RAD21 or NIPBL. Red arrows indicate lagging chromosomes. Right, quantification of cells with sharply separated DNA. Data are presented as mean ± SEM. Two-tailed unpaired Student’s t-test; **P < 0.01, ns, not significant.

**Figure S2.**
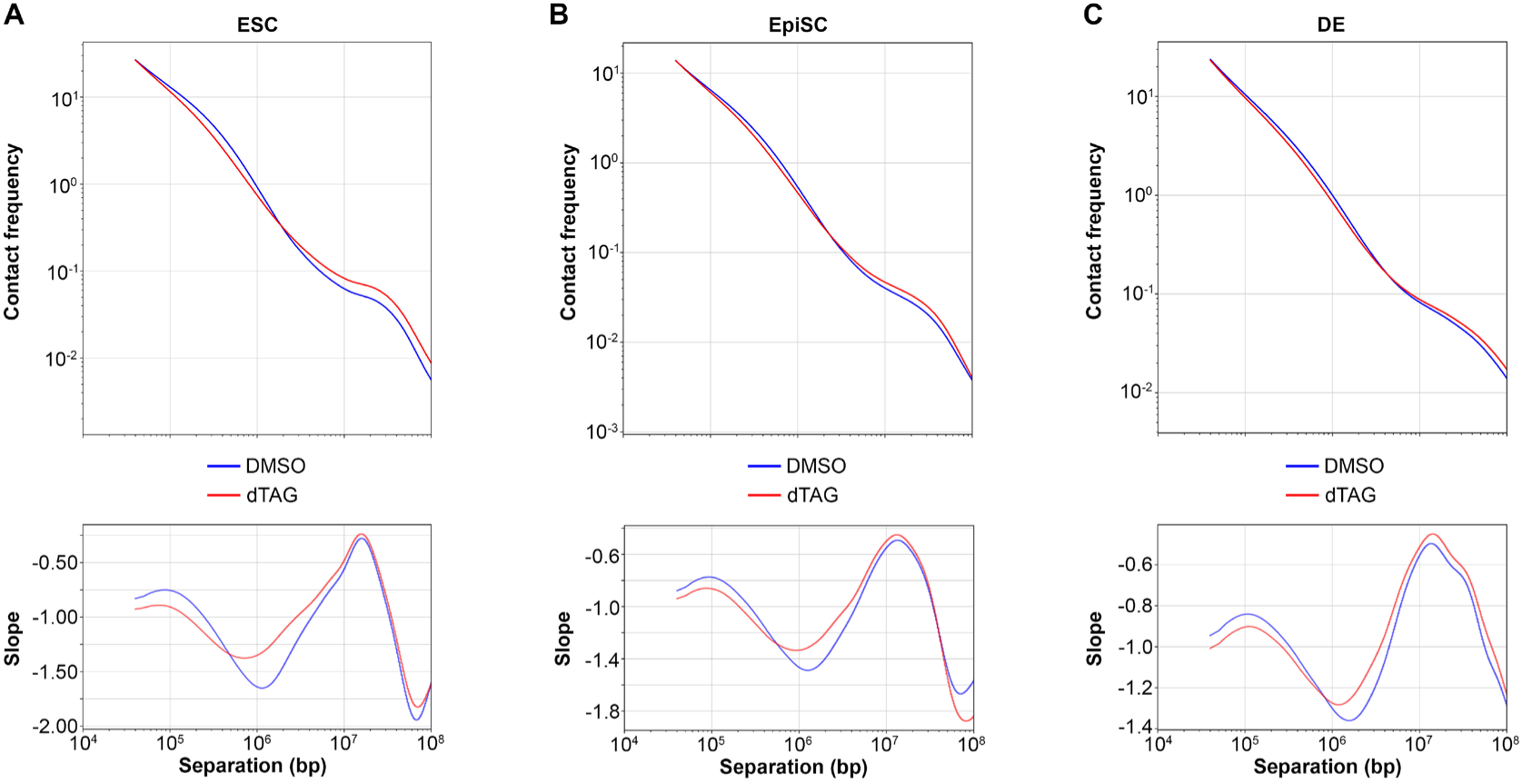
Altered chromatin contact frequencies upon NIPBL depletion in mESCs, EpiSCs and definitive endoderm. **(A)** Genome-wide contact probability and slope curves from Hi-C data of 4 h DMSO- or dTAG-treated mouse ESCs (10 kb bin size). **(B)** Genome-wide contact probability and slope curves from Hi-C data of 4 h DMSO- or dTAG-treated EpiSCs (10 kb bin size). **(C)** Genome-wide contact probability and slope curves from Hi-C data of DMSO- or dTAG-treated mouse DE cells (10 kb bin size). Details of the dTAG treatment time course are presented in Fig. 5I and described in the Methods section.

**Figure S3.**
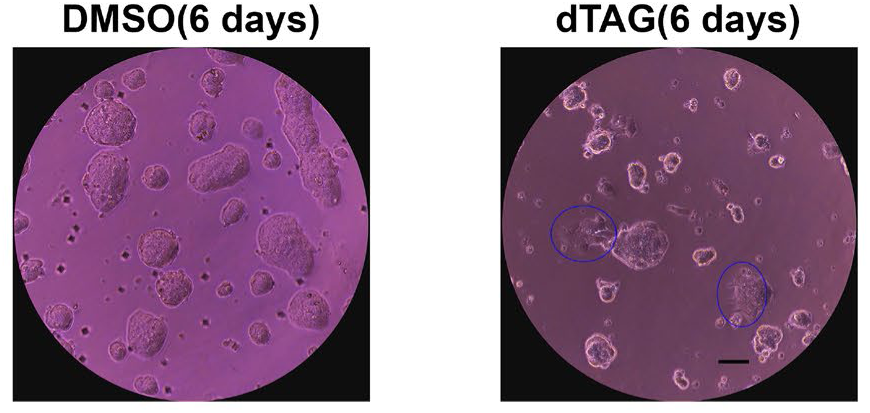
Brightfield images showing mESC colonies after treatment with DMSO or dTAG for 6 days. Blue circles highlight differentiated cells in the dTAG-treated sample. Scale bar = 100 μm.

**Figure S4.**
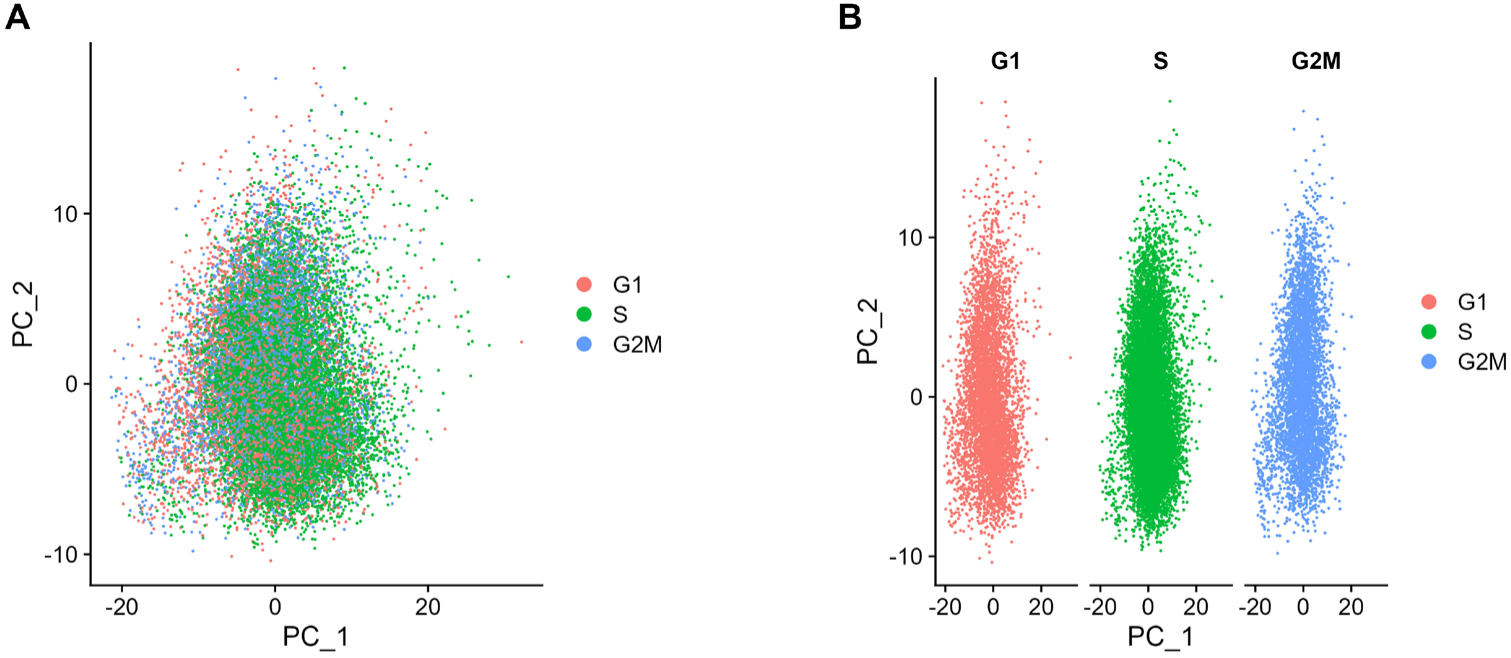
Cell cycle phase analysis showed that cells in different phases were evenly distributed across clusters. (A)(B) Principal component analysis (PCA) of scRNA-seq data from mouse ESCs colored by cell-cycle phases (G1, S, and G2/M).

**Figure S5.**
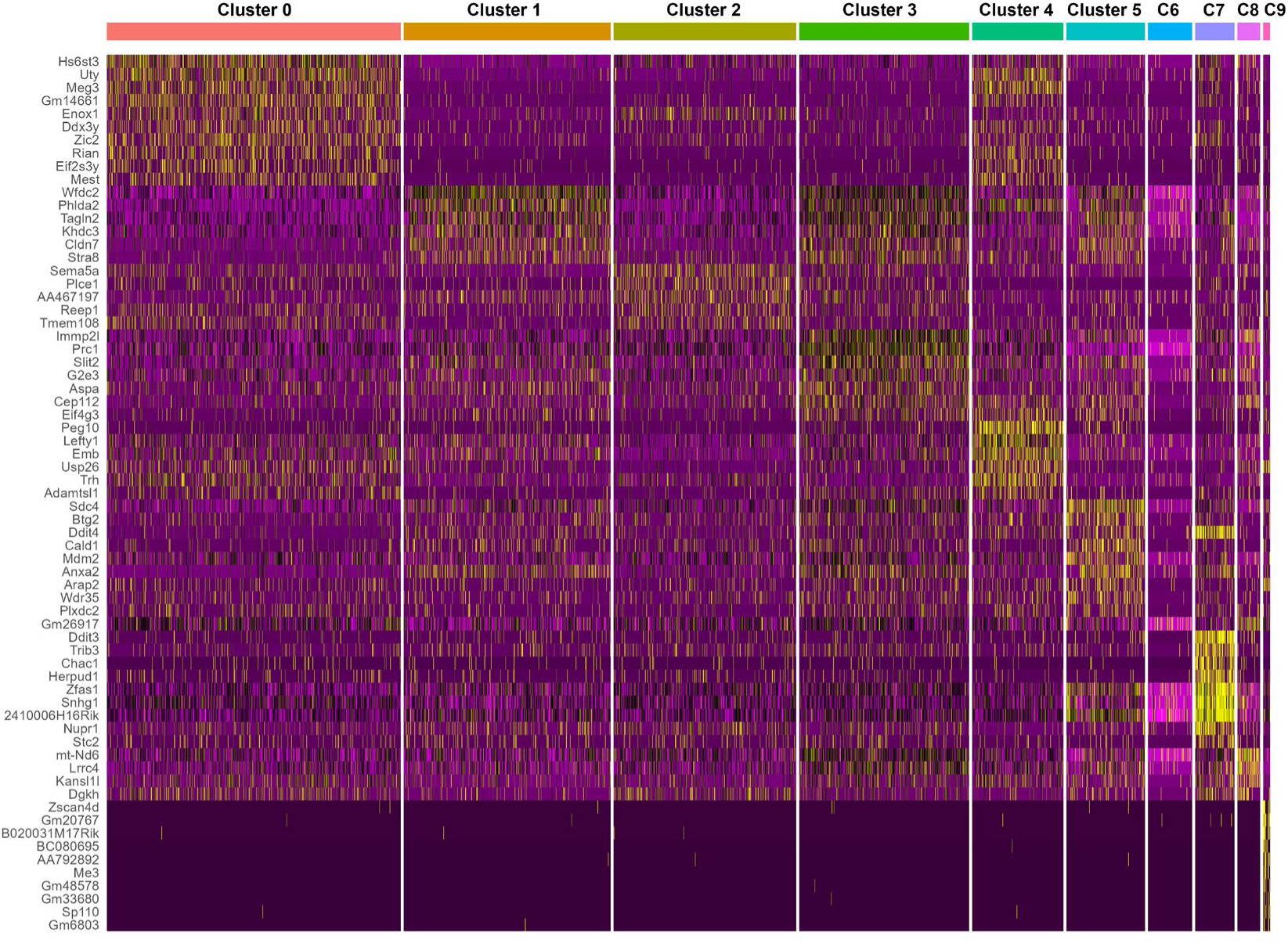
Top differentially expressed genes in each cluster identified from scRNA-seq data of mouse ESCs.

**Figure S6.**
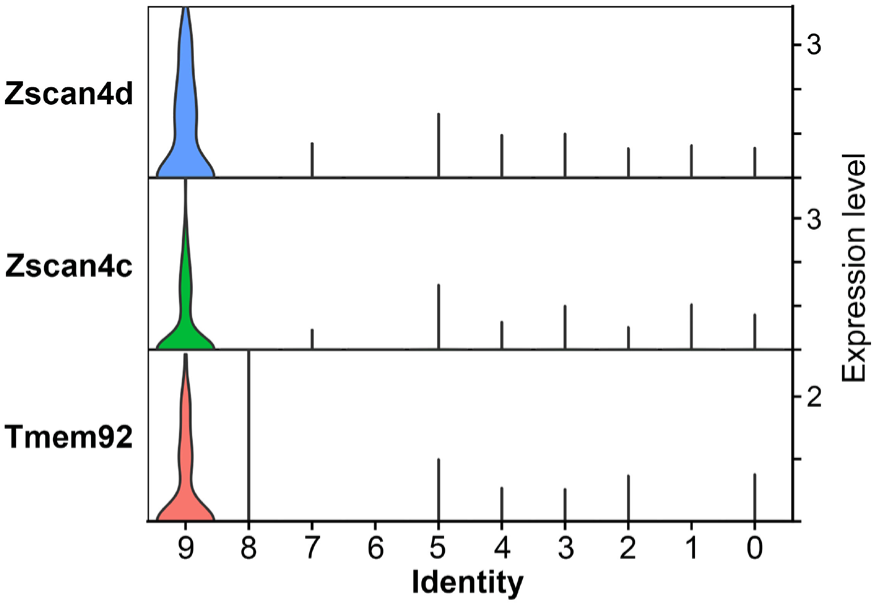
Violin plots illustrating the expression levels of 2-cell embryo markers across cell clusters.

**Figure S7.**
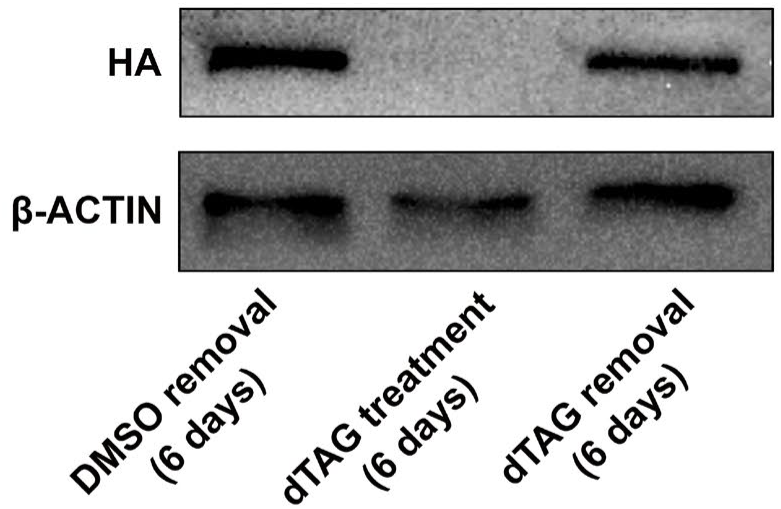
Western blot analysis of HA–NIPBL in mESCs treated with DMSO or dTAG for 6 days, followed by dTAG removal for an additional 6 days. β-ACTIN was used as a loading control.

**Figure S8.**
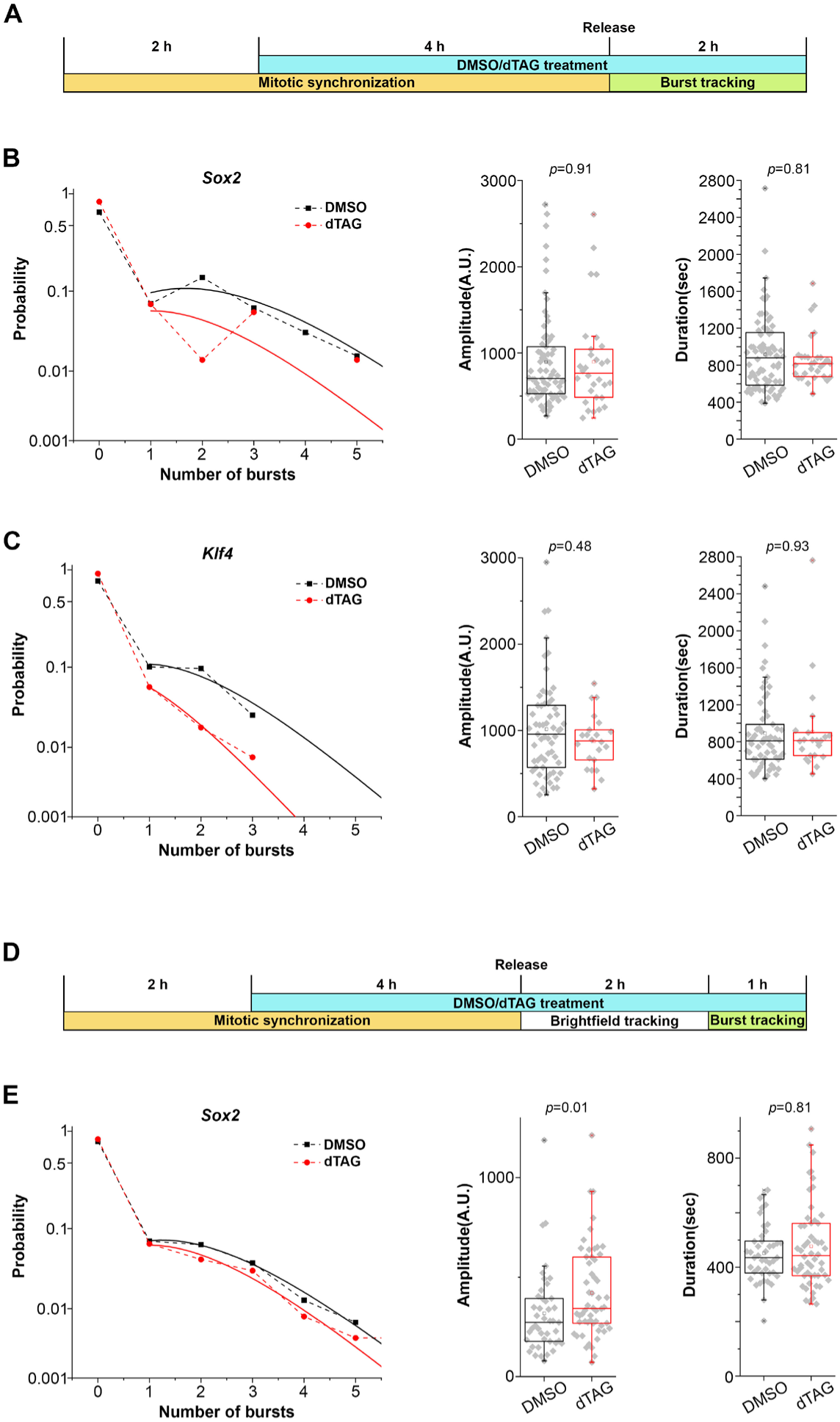
*Sox2*, but not *Klf4*, reactivation during cell division is more dependent on NIPBL than the maintenance of its expression afterwards. **(A)** Schematic overview of *Sox2* and *Klf4* reactivation experiments. **(B)(C)** Burst probability, amplitude, and duration of *Sox2* and *Klf4* during reactivation in DMSO- and dTAG-treated cells. Each gray dot in the box plots of burst amplitude and duration represents one burst. Data are from two independent experiments. **(D)** Schematic overview of post-reactivation *Sox2* experiments. **(E)** Burst probability, amplitude, and duration of *Sox2* post-reactivation in DMSO- and dTAG-treated cells. Each gray dot in the box plots of burst amplitude and duration represents one burst. Data are from two independent experiments.

**Figure S9.**
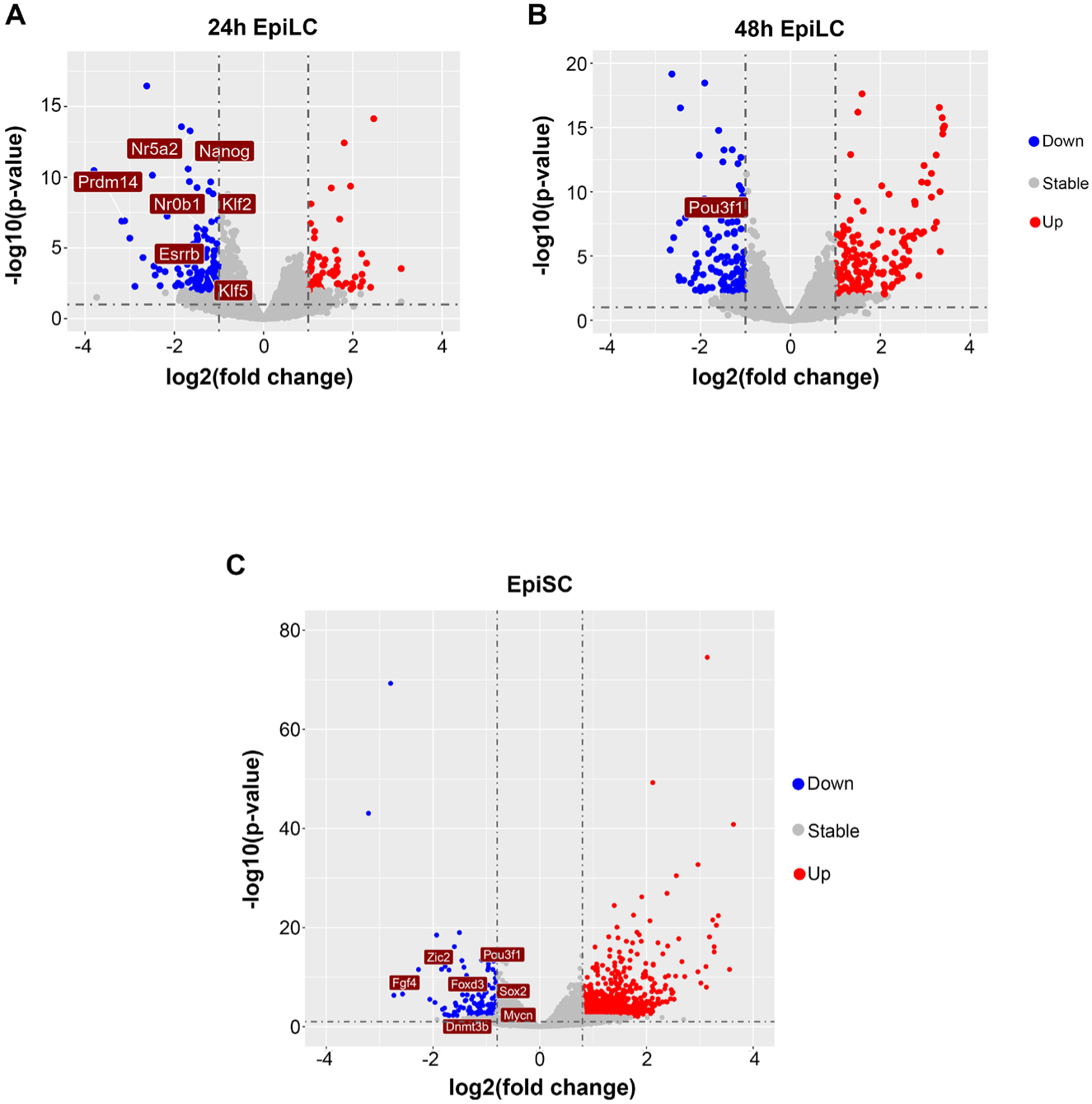
Differentially expressed genes in EpiLCs and EpiSCs after NIPBL depletion. **(A)(B)(C)** Volcano plots showing differentially expressed genes between DMSO- and dTAG-treated EpiLCs and EpiSCs, following the treatment scheme shown in Fig. 5A-C. Upregulated and downregulated genes are shown in red and blue, respectively, while genes without significant changes are shown in gray (P < 0.01; |fold change| ≥ 2). Labeled genes represent downregulated genes in the corresponding samples.

**Figure S10.**
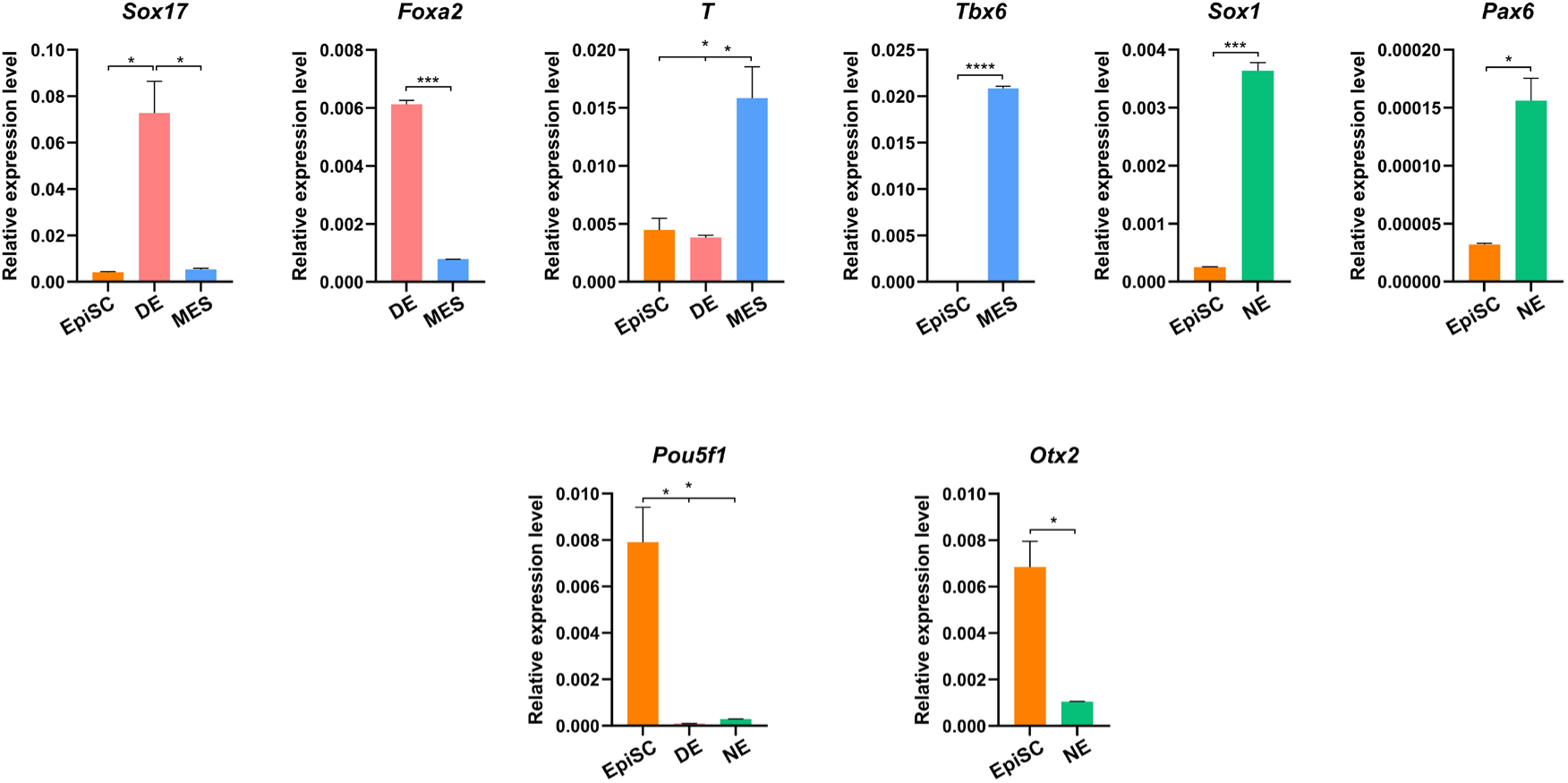
qPCR analysis of marker gene expression for the three germ layers: *Sox17* and *Foxa2* for definitive endoderm (DE), *T* and *Tbx6* for mesoderm (MES), and *Sox1* and *Pax6* for neuroectoderm (NE). *Pou5f1* showed low expression levels in DE and NE, while *Otx2* showed low expression in NE. Values were normalized to *Gapdh* expression. Data are presented as mean ± SEM. Two-tailed unpaired Student’s t-test; *P<0.05, **P < 0.01, ***P < 0.001, ****P<0.0001.

**Figure S11.**
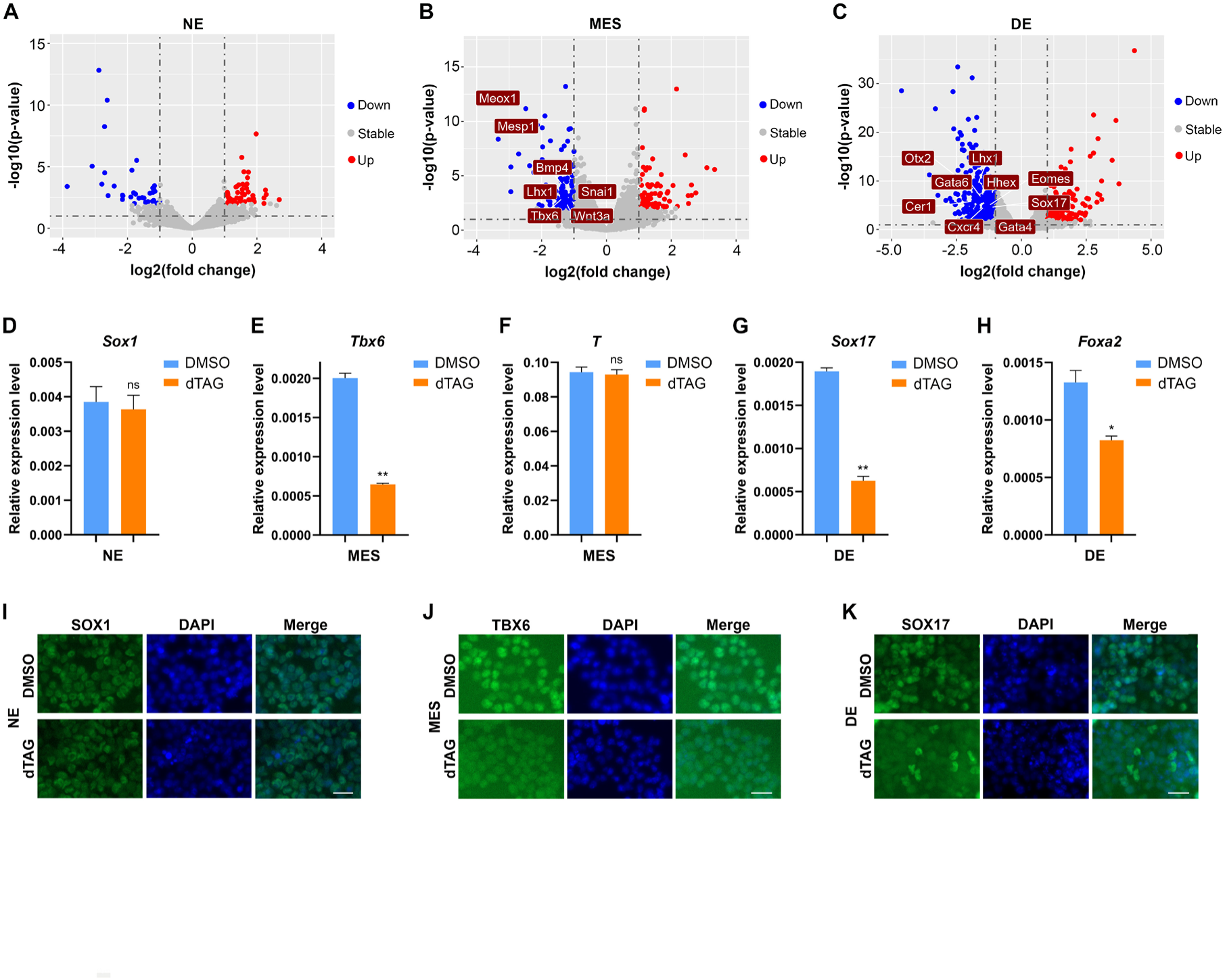
Distinct effects of NIPBL depletion on the formation of the three germ layers. **(A)(B)(C)** Volcano plots showing differentially expressed genes between DMSO- and dTAG-treated neuroectoderm (NE), mesoderm (MES), and definitive endoderm (DE). Upregulated and downregulated genes are shown in red and blue, respectively, while genes without significant changes are shown in gray (P < 0.01; |fold change| ≥ 2). Labeled genes represent downregulated genes in the corresponding samples. **(D)(E)(F)(G)(H)** qPCR analysis of marker gene expression for the three germ layers after treatment with DMSO or dTAG. Values were normalized to *Gapdh* expression. Data are presented as mean ± SEM. Two-tailed unpaired Student’s t-test; *P < 0.05, **P < 0.01, ns, not significant. **(I)(J)(K)** Immunofluorescence staining of SOX1, TBX6, and SOX17 in the three germ layers after treatment with DMSO or dTAG. Scale bar: 20 μm. All experiments are performed following the treatment scheme shown in Fig. 5G–I.

**Figure S12.**
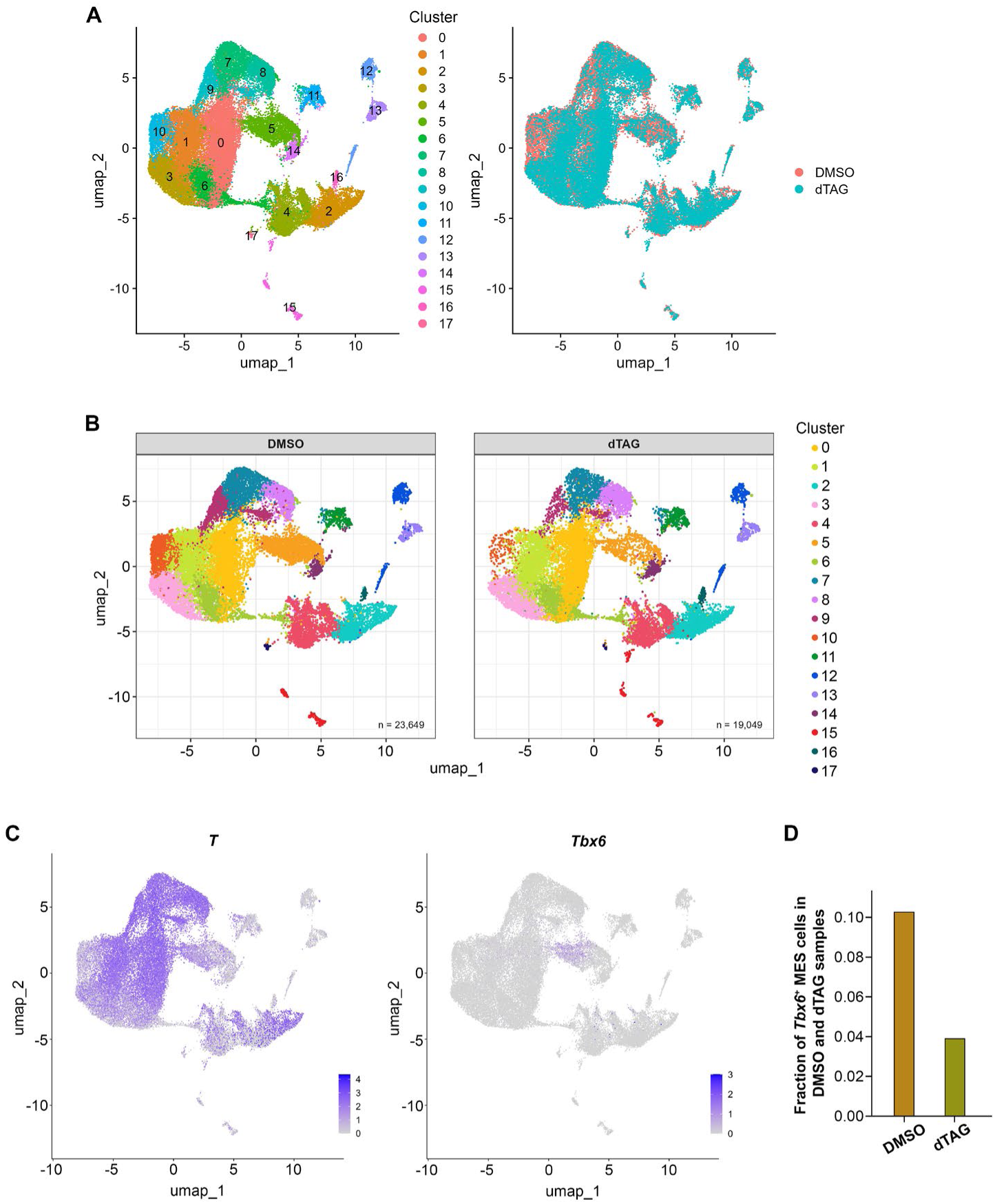
Decreased proportion of *Tbx6*-positive mesodermal cells in the differentiated population. **(A)** Left: UMAP plot showing cell clusters from the integrated datasets of mesoderm cells treated with DMSO or dTAG, following the treatment scheme shown in Fig. 5H. Right: UMAP plot showing DMSO-treated mesoderm cells in red and dTAG-treated mesoderm cells in blue. **(B)** UMAP plots split by sample. **(C)** Feature plots showing the expression of *T* and *Tbx6* in each cell cluster. Cluster 5 represents *Tbx6*-positive mesoderm. Bar graph (right): Fraction of *Tbx6*-positive MES cells in DMSO- and dTAG-treated samples.

**Figure S13.**
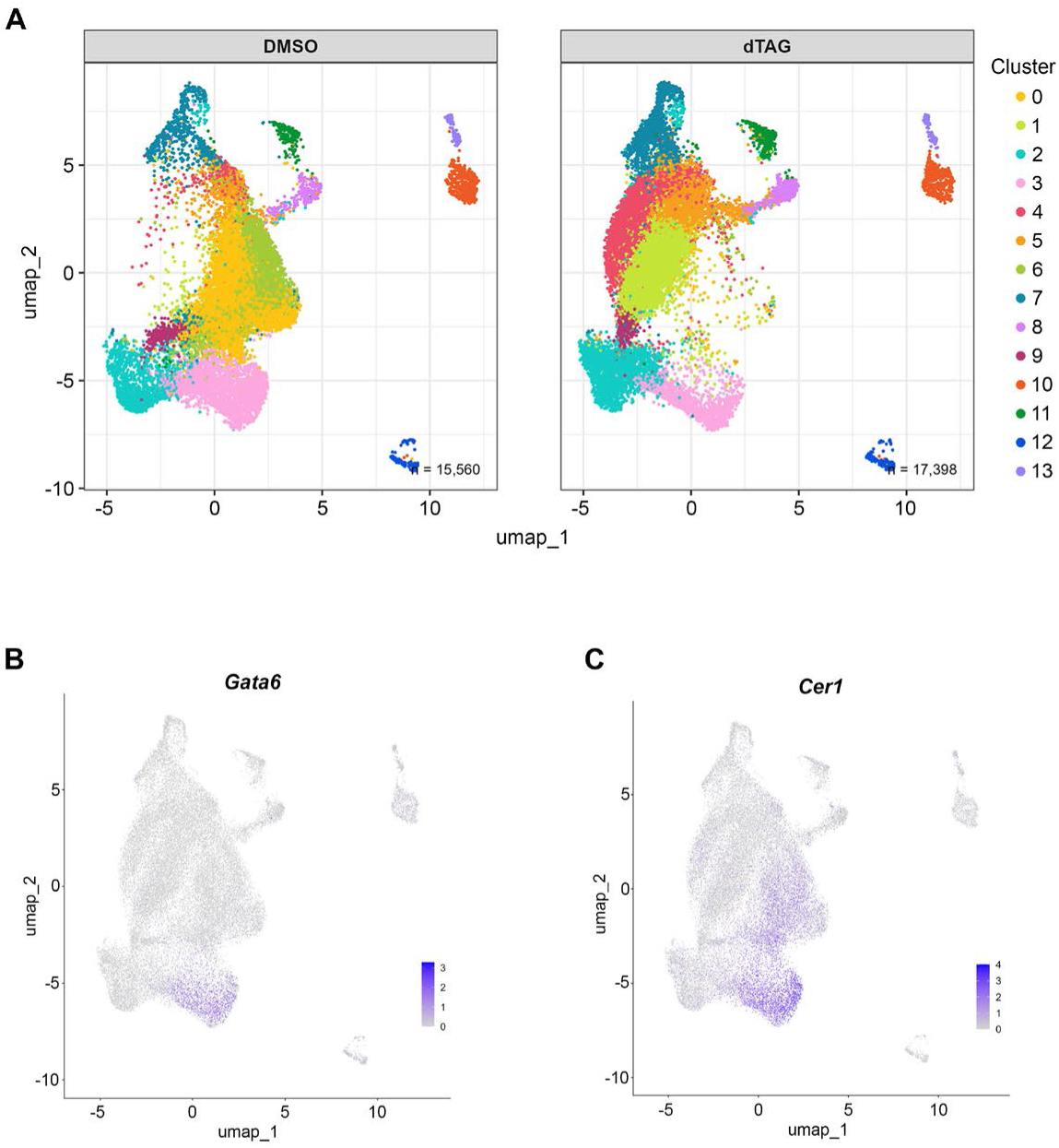
NIPBL depletion results in abnormal PS-like cells and consequently compromises definitive endoderm formation. **(A)** UMAP plots split by DMSO- or dTAG-treated definitive endoderm samples, showing the distribution of cells under each condition. **(B)** Feature plots showing the expression of *Gata6* and *Cer1* in each cell cluster.

**Figure S14.**
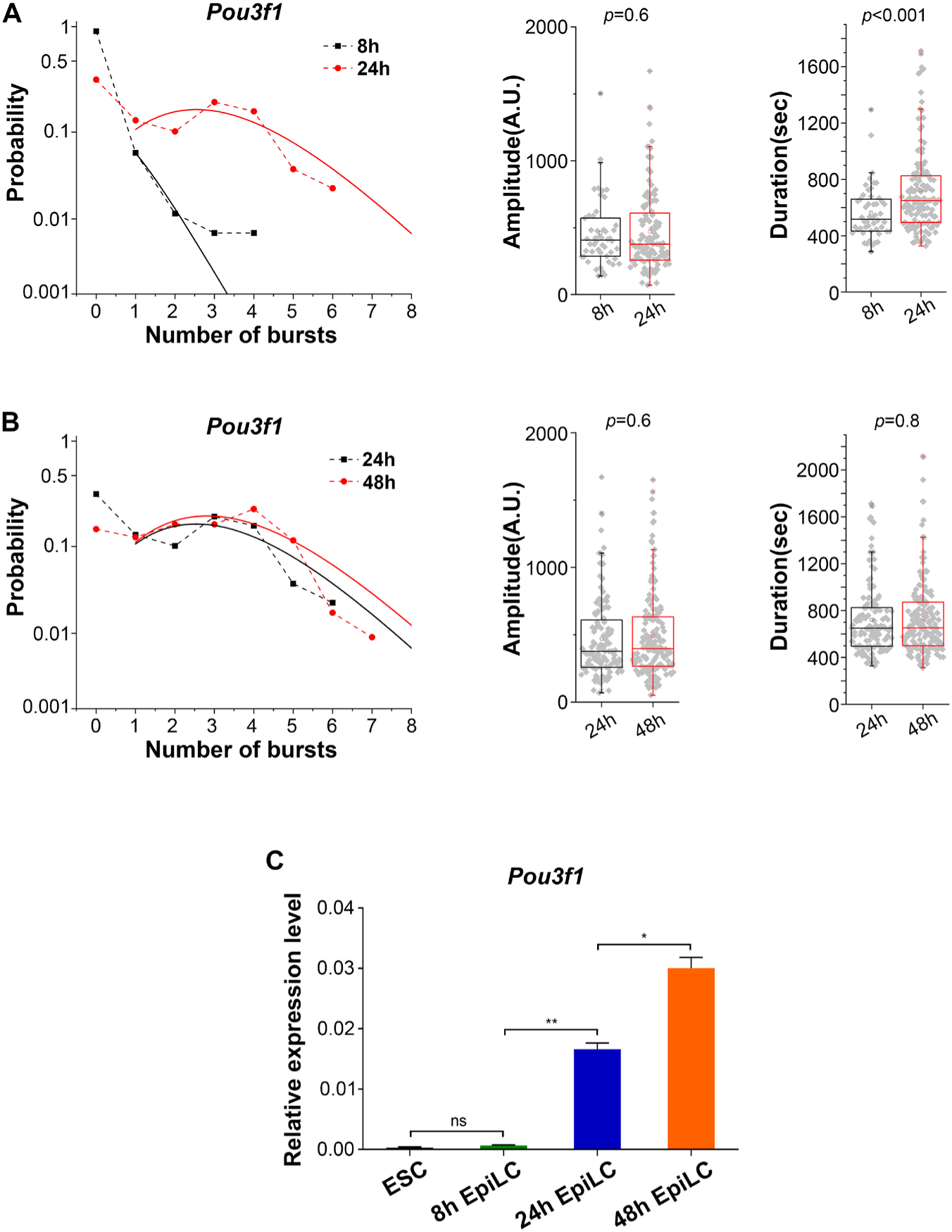
The dynamics of *de novo Pou3f1* activation during EpiLC differentiation. **(A)** Burst probability, amplitude, and duration of *Pou3f1* in 8h and 24h EpiLCs. Each gray dot in the box plots of burst amplitude and duration represents one burst. Data are from two independent experiments. **(B)** Burst probability, amplitude, and duration of *Pou3f1* in 24h and 48h EpiLCs. Each gray dot in the box plots of burst amplitude and duration represents one burst. Data are from two independent experiments. **(C)** qPCR analysis of *Pou3f1* expression during EpiLC differentiation. Values were normalized to *Gapdh* expression. Data are presented as mean ± SEM. Two-tailed unpaired Student’s t-test; *P<0.05, **P < 0.01, ns, not significant.

**Figure S15.**
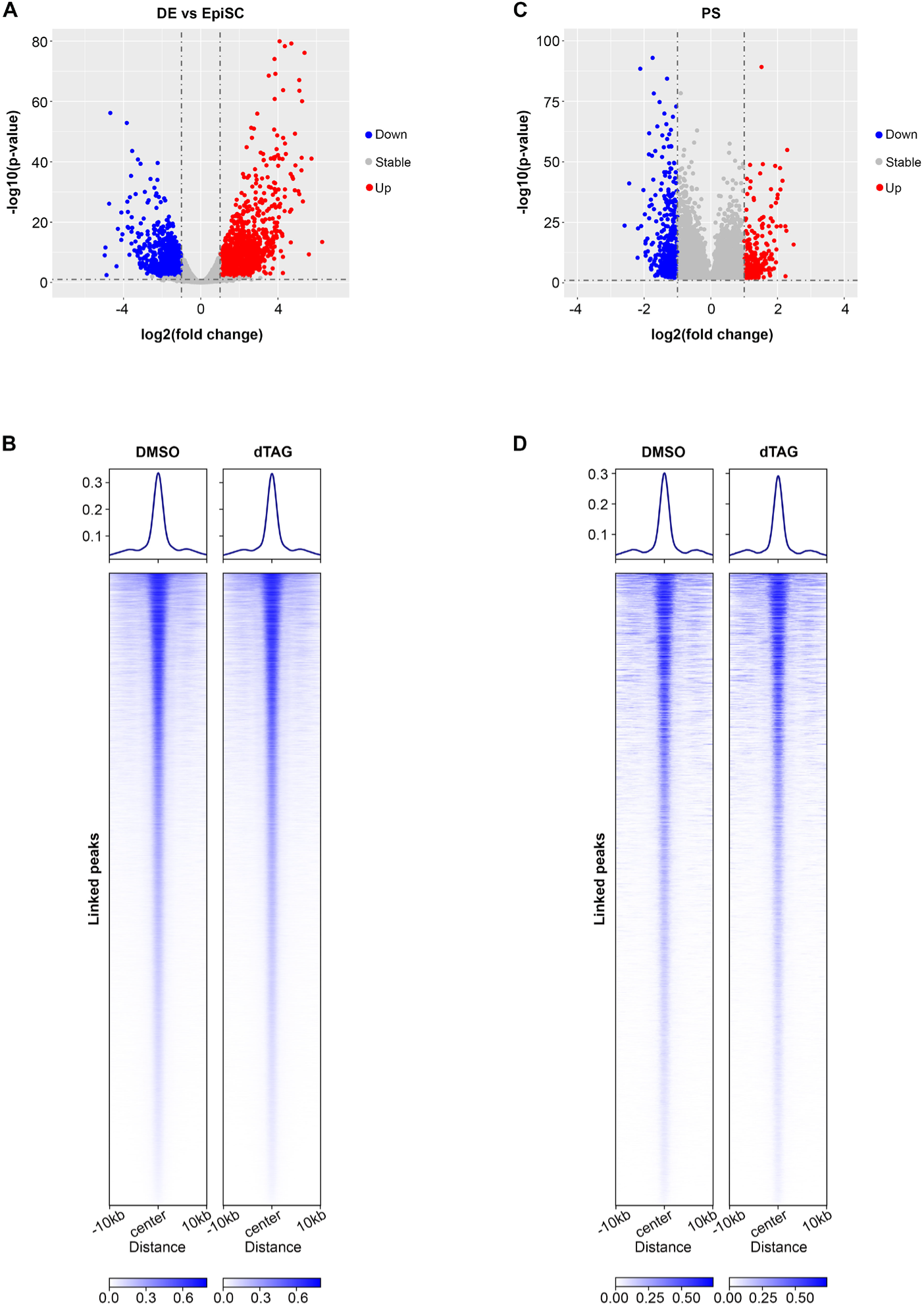
There are no obvious changes in chromatin accessibility of likely cis-elements caused by NIPBL depletion in PS-like cells. **(A)** Volcano plots showing differentially expressed genes between EpiSC and definitive endoderm (DE). Upregulated and downregulated genes in DE are shown in red and blue, respectively, while genes without significant changes are shown in gray (P < 0.01; |fold change| ≥ 2). **(B)** Heatmaps and profiles showing subtle changes in chromatin accessibility of the activated and deactivated genes highlighted in (**A**) in DE cells after NIPBL depletion. The Peak2GeneLinkage function in ArchR was used to identify peaks associated with activated or deactivated genes based on scMultiome data. *Col1a1*⁺ feeder cells were excluded (Clusters 10, 12, and 13 in Fig. 5M). **(C)** Volcano plots showing differentially expressed genes (DEGs) that overlap between the activated/deactivated genes highlighted in (**A**) and DEGs identified from primitive streak (PS) clusters after NIPBL depletion based on scMultiome data of DE cells. Upregulated and downregulated genes are shown in red and blue, respectively, while genes without significant changes are shown in gray (P < 0.01; |fold change| ≥ 2). **(D)** Heatmaps and profiles showing subtle changes in chromatin accessibility of DEGs highlighted in (**C**) after NIPBL depletion. The Peak2GeneLinkage function in ArchR was used to identify peaks associated with DEGs based on scMultiome data.

**Figure S16.**
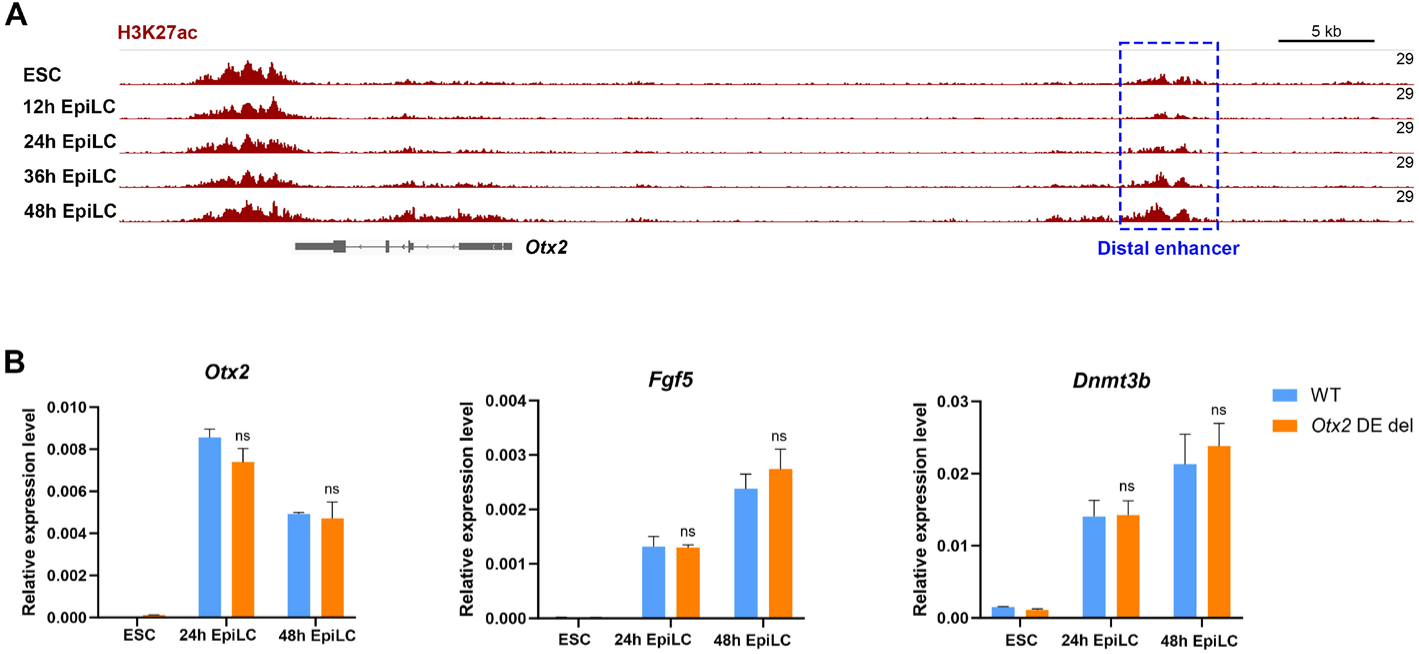
Deletion of the distal enhancer does not affect activation of *Otx2* or other EpiLC markers. **(A)** ChIP-seq signal tracks of H3K27ac around the *Otx2* locus during the time course of mESC differentiation into EpiLCs, using published datasets. Distal enhancers are highlighted by blue dashed box. **(B)** qPCR analysis of *Otx2*, *Fgf5*, and *Dnmt3b* expression during the differentiation of mESCs into EpiLCs before and after distal enhancer deletion. Values were normalized to *Gapdh* expression. Data are presented as mean ± SEM. Two-tailed unpaired Student’s t-test; ns, not significant.

**Figure S17.**
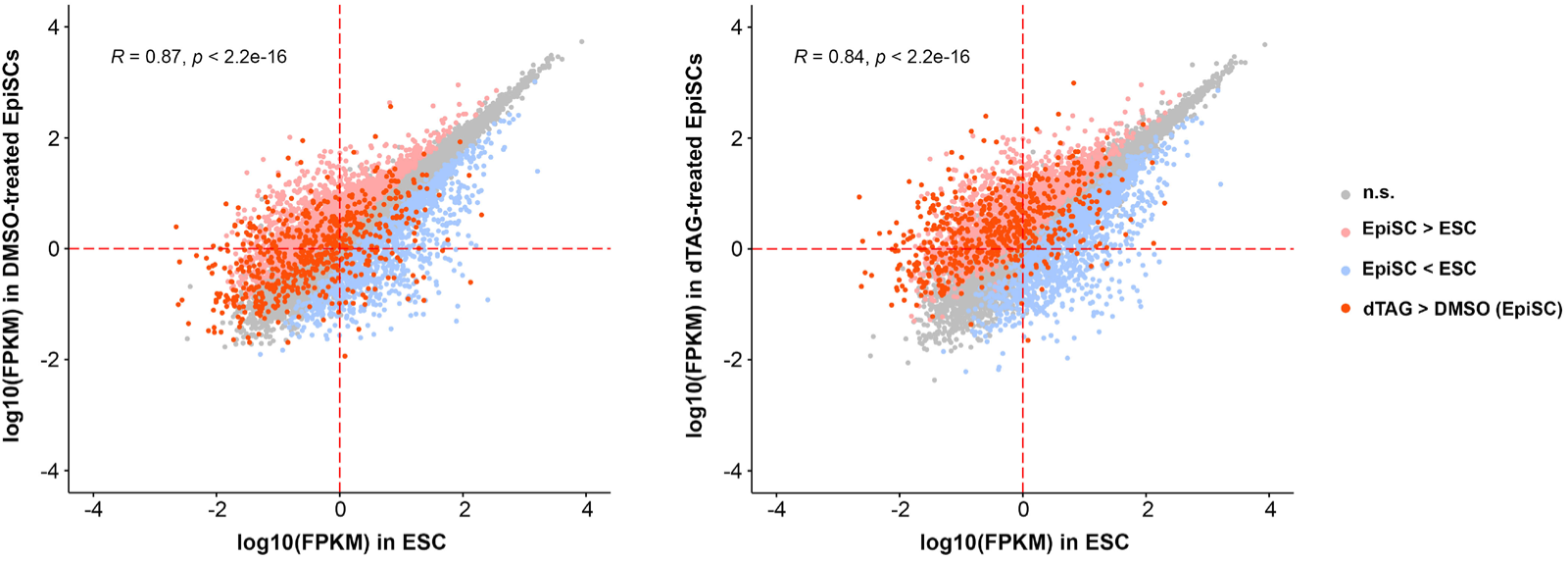
Long-term NIPBL depletion upregulates repressed genes. Correlation plot of gene expression levels (log₁₀ FPKM) between ESCs and DMSO- or dTAG-treated EpiSCs from an independent biological replicate. Upregulated and downregulated genes in EpiSCs are shown in pink and light blue, respectively, while genes without significant changes are shown in gray. Genes upregulated upon NIPBL depletion in EpiSCs are shown in red. P < 0.01; |fold change| ≥ 2.

**Figure S18.**
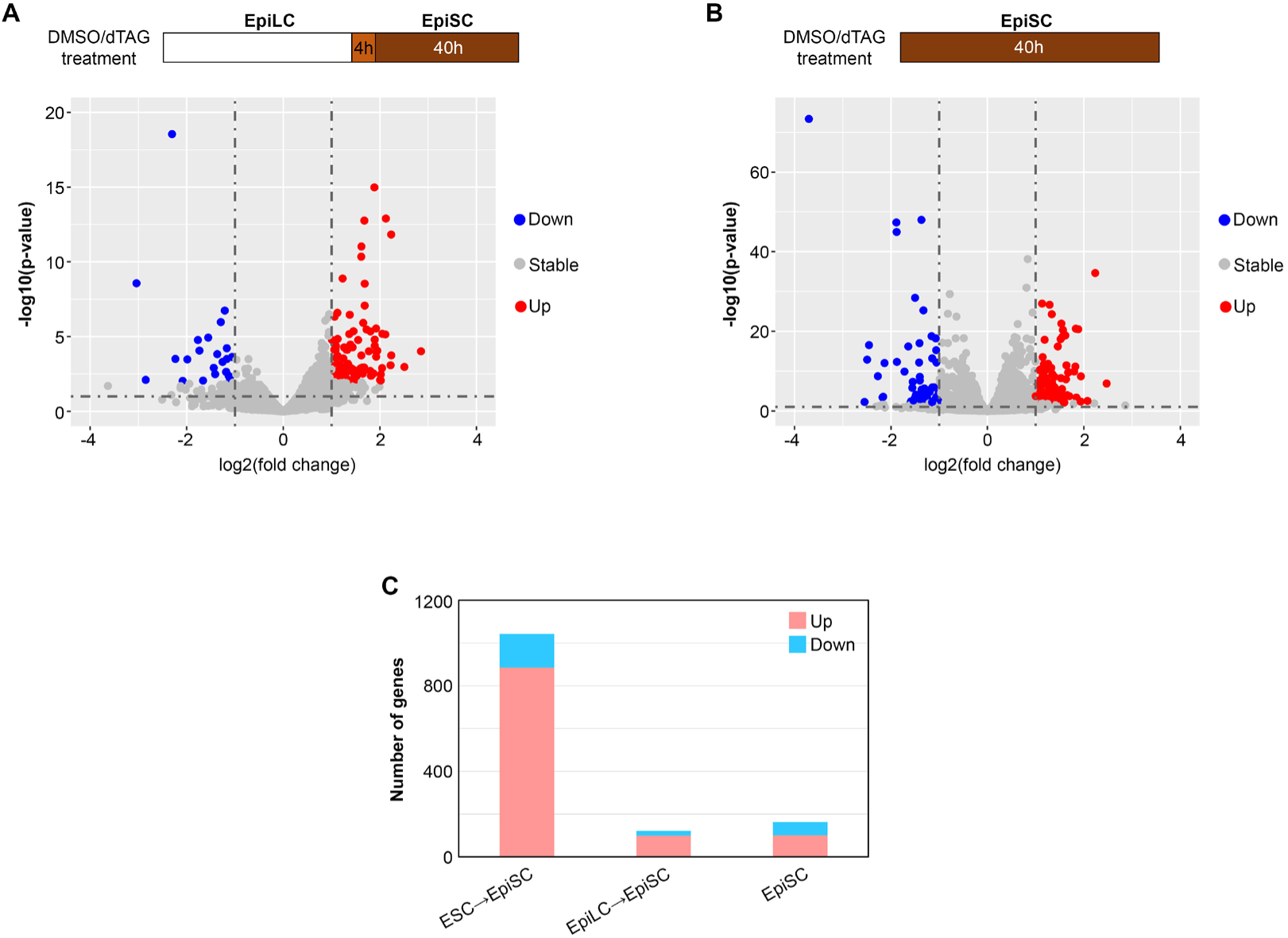
Aberrant gene upregulation is not specific to the final EpiSC state or intermediate EpiLC-to-EpiSC transition. **(A)(B)** Upper: Schematic overview of the timing of DMSO or dTAG treatment in EpiSCs and during the EpiLC-to-EpiSC transition. Lower: Volcano plots showing differentially expressed genes between DMSO- and dTAG-treated EpiSCs. Upregulated and downregulated genes are shown in red and blue, respectively, while genes without significant changes are shown in gray (P < 0.01; |fold change| ≥ 2). **(C)** Bar graph showing the number of upregulated and downregulated genes across three experiments. ESC-to-EpiSC experiments follow the treatment scheme shown in Fig. 5C. EpiLC-to-EpiSC transition and EpiSC experiments follow the treatment schemes shown in the upper panels of (**A**) and (**B**).

**Figure S19.**
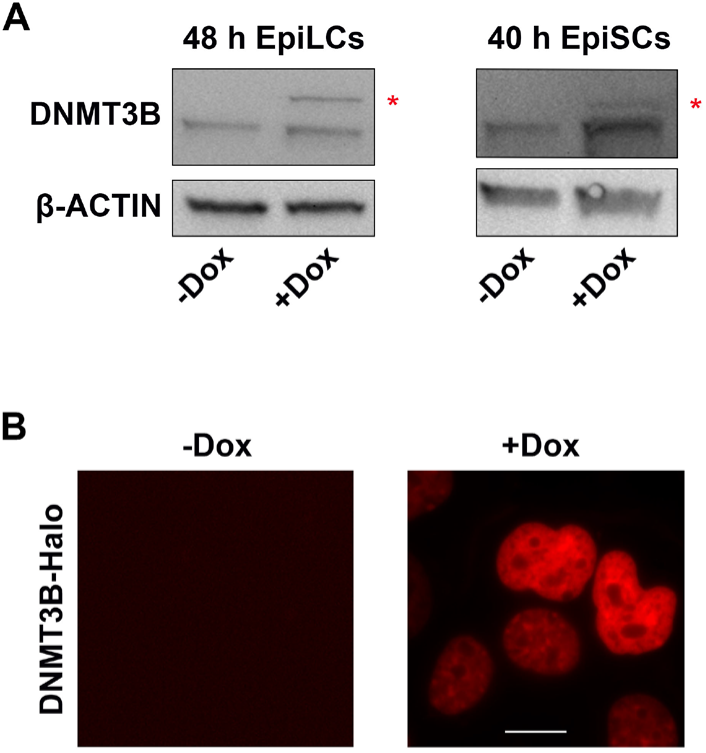
The efficiency of DNMT3B overexpression. **(A)** Western blot analysis of DNMT3B in EpiLCs and EpiSCs followed the treatment scheme shown in Fig. 8d. β-ACTIN was used as a loading control. Red asterisks indicate the fused DNMT3B-Halo protein bands. **(B)** Representative fluorescence images showing overexpression of DNMT3B-Halo in 48 h EpiLCs. DNMT3B-Halo is labeled with HaloTag ligand prior to imaging. Scale bar = 10 µm.

**Figure S20.**
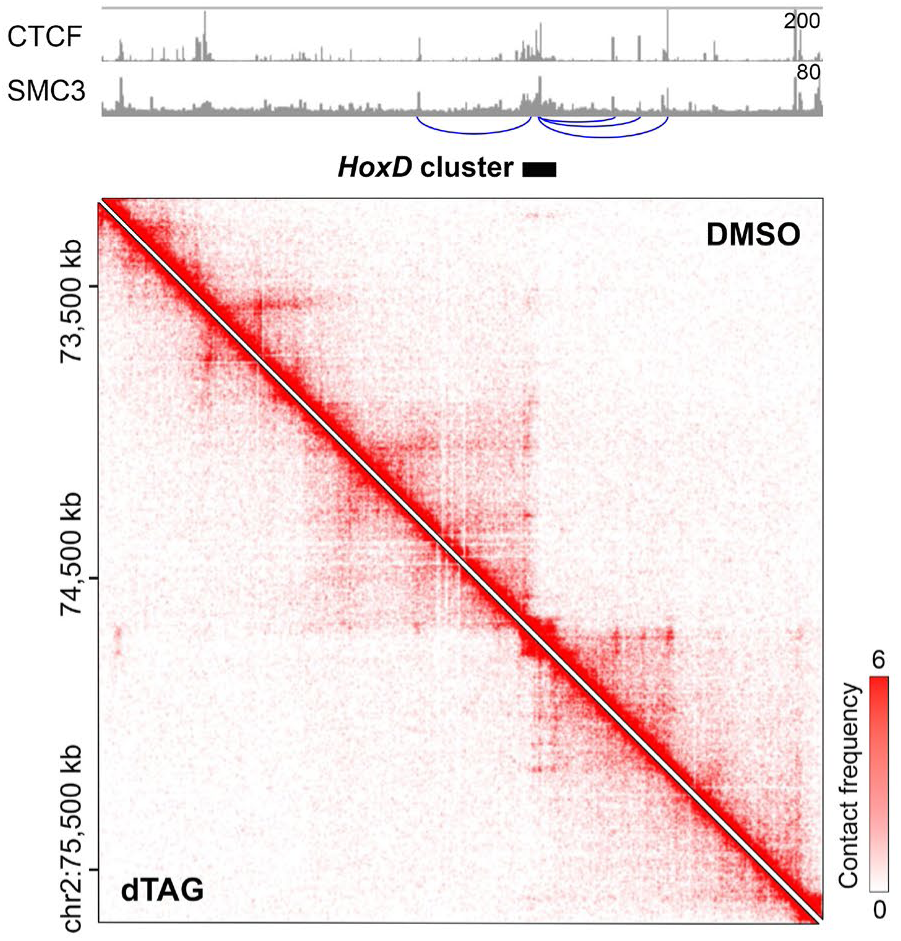
Hi-C snapshots showing chromatin contacts in DMSO-treated (top right) and dTAG-treated (bottom left) EpiSC around the *HoxD* cluster. Top panels show ChIP-seq signal tracks for CTCF and SMC3 at the same genomic regions, using published datasets from E6.5 mouse embryos.

## Notes

### Competing Interest Statement

The authors have declared no competing interest.

